# Mitophagy protects beta cells from inflammatory damage in diabetes

**DOI:** 10.1101/2020.06.07.138917

**Authors:** Vaibhav Sidarala, Gemma L. Pearson, Vishal S. Parekh, Benjamin Thompson, Lisa Christen, Morgan A. Gingerich, Jie Zhu, Tracy Stromer, Jianhua Ren, Emma Reck, Biaoxin Chai, John A. Corbett, Thomas Mandrup-Poulsen, Leslie S. Satin, Scott A. Soleimanpour

**Affiliations:** Division of Metabolism, Endocrinology & Diabetes and Department of Internal Medicine, University of Michigan Medical School, Ann Arbor, MI 48105, USA; Department of Pharmacology, University of Michigan Medical School, Ann Arbor, MI 48105, USA; Department of Biomedical Sciences, University of Copenhagen, Copenhagen, Denmark; Program in Biological Sciences, University of Michigan Medical School, Ann Arbor, MI 48105, USA; Department of Biochemistry, Medical College of Wisconsin, Milwaukee, WI 53226, USA; VA Ann Arbor Healthcare System, Ann Arbor, MI 48105, USA

## Abstract

Inflammatory damage contributes to β-cell failure in type 1 and 2 diabetes (T1D and T2D). Mitochondria are damaged by inflammatory signaling in β-cells, resulting in impaired bioenergetics and initiation of pro-apoptotic machinery. Hence, the identification of protective responses to inflammation could lead to new therapeutic targets. Here we report that mitophagy serves as a protective response to inflammatory stress in both human and rodent β-cells. Utilizing *in vivo* mitophagy reporters, we observed that diabetogenic pro-inflammatory cytokines induced mitophagy in response to nitrosative/oxidative mitochondrial damage. Mitophagy-deficient β-cells were sensitized to inflammatory stress, leading to the accumulation of fragmented dysfunctional mitochondria, increased β-cell death, and hyperglycemia. Overexpression of *CLEC16A*, a T1D gene and mitophagy regulator whose expression in islets is protective against T1D, ameliorated cytokine-induced human β-cell apoptosis. Thus, mitophagy promotes β-cell survival and prevents diabetes by countering inflammatory injury. Targeting this pathway has the potential to prevent β-cell failure in diabetes and may be beneficial in other inflammatory conditions.

---

Inflammatory stress plays a crucial role in the pathogenesis of several metabolic diseases, including diabetes (1-3). All forms of diabetes share a common etiology of insufficient pancreatic β-cell mass or function to meet peripheral insulin demand, and inflammatory injury is commonly associated with β-cell dysfunction (1, 4, 5). Although the precise molecular mechanisms are unclear, the excess generation of free radicals, including nitric oxide (NO) and/or reactive oxygen species (ROS), likely contribute to β-cell inflammatory damage (6). Mitochondria are adversely affected by inflammatory signaling in β-cells, which can result in impaired bioenergetics, blunted glucose-stimulated insulin secretion, and activation of apoptosis (7). Therefore, strategies to block inflammation and/or preserve mitochondrial function are of great interest as potential diabetes therapies.

Mitochondria exist in dynamic networks that undergo continuous remodeling and respond to both internal and external environmental cues. Mitophagy is an essential quality control mechanism that selectively eliminates damaged mitochondria to maintain healthy mitochondrial networks (8-10). The T1D susceptibility gene *CLEC16A* encodes an E3 ubiquitin ligase that controls mitophagic flux in β-cells (11-13), indicating a critical role for mitophagy in maintaining β-cell function. Indeed, diabetogenic intronic polymorphisms in the *CLEC16A* locus that reduce human islet CLEC16A mRNA expression are associated with impaired β-cell function and glucose control in humans (13, 14). Although mitophagy maintains the metabolic function needed for glucose-stimulated insulin release, it has not been shown to affect β-cell survival (11, 13, 15). Further, whether mitophagy (or Clec16a) protects β-cells from inflammatory attack is unknown.

Here we elucidate a key protective role for mitophagy in the response to inflammatory stress in β-cells. Utilizing *in vivo* mitochondrial biosensors and biochemical/genetic approaches, we show that pro-inflammatory cytokines, which model the inflammation that occurs during diabetes pathogenesis, induce mitophagy in both human and rodent β-cells. Cytokine-induced free radicals function as upstream inflammatory signals to activate β-cell mitophagy, and the impairment of Clec16a-mediated mitophagy exacerbates hyperglycemia and β-cell apoptosis following inflammatory stimuli. Lastly, we demonstrate that adenoviral overexpression of CLEC16A protects human β-cells against cytokine-mediated demise, illustrating the feasibility of therapeutically targeting this process.

## Materials and methods

### Animals

Animal studies were approved by the University of Michigan Institutional Animal Care and Use Committee. Mouse models included *Clec16a*^loxP^ (13), *Ins1*-Cre (16), mtKeima ((17); a gift from Toren Finkel), and *NOS2*^-/-^ mice (18). mtKeima mice were separately maintained on the FVB/N and C57BL/6N (B6N) backgrounds, while all other models were maintained on the B6N background. *Clec16a*^loxP^ were mated to *Ins1*-Cre mice, and B6N mtKeima were mated to *NOS2*^-/-^ mice to generate experimental groups. For studies using the conditional Clec16a allele, *Ins1*-Cre alone mice and *Clec16a*^loxP/loxP^ mice were phenotypically indistinguishable from each other and thus used as “WT” controls. For streptozotocin (STZ) studies, 5-week-old male mice were injected with 50 mg/kg STZ (Cayman Chemical) for five consecutive days (Day 1-5). Random fed blood glucose concentrations were measured 3-4 times weekly for up to 35 days post-STZ. Animals were housed on a standard 12hr light/12hr dark cycle with *ad libitum* access to food and water.

### Islet isolation and cell culture

Islets were isolated from male and female mice between the ages of 8-15 weeks and cultured as previously described (11, 13). Min6 β-cells (passages 28-40) were maintained as previously described (11, 13). Human islets were acquired from the Integrated Islet Distribution Program (IIDP) and cultured as previously described (11). Donor information is provided in Supplementary Table 1.

### Cell treatments, transfections, and viral transductions

Min6 β-cells and mouse islets were exposed to a cytokine combination consisting of murine IL-1β, TNF-α, and IFN-γ (Peprotech) for 6-24hrs. Human islets were treated with a cytokine mixture containing human IL-1β, TNF-α, and IFN-γ (Peprotech) for 24-48hrs. Additional cell treatments included dimethylsulfoxide as a vehicle control (DMSO; Fisher), lenalidomide (Sigma), tolbutamide (Sigma), FCCP (Sigma), valinomycin (Sigma), rotenone (Sigma), paraquat (Sigma), DPTA/NO (Cayman Chem), L-NMMA (Cayman Chem), and tiron (Sigma). Min6 β-cells were transfected using an Amaxa Nucleofector (Lonza) as previously described (13). The tandem mitochondrial-targeted mCherry-eGFP mitophagy reporter plasmid (pCLBW-cox8-EGFP-mCherry) was a gift from David Chan (Addgene 78520). pLKO.1-shNT, pLKO.1-shClec16a, pFLAG-CMV5a-empty vector, and pFLAG-CMV5a-Clec16a constructs were previously described (11, 13) and used for knockdown or overexpression studies. Adenoviral vectors and particles were generated (VectorBuilder) expressing Clec16a-3xFLAG (Ad.Clec16a) or an empty vector control (Ad.EV), as well as an IRES:GFP, which were driven by the 2.9kb Pdx1 proximal promoter sequence (19) for β-cell-selective expression. Intact human islets (100 IEQs) were transduced with 1×10^9^ PFU of Ad.Clec16a or Ad.EV for 24hrs, prior to cytokine exposure 48hrs post-infection for cell survival studies. β-cell transduction was confirmed by insulin and GFP co-staining. Human islets were transduced overnight with adenoviral particles expressing the Perceval-HR biosensor (described below) 48hrs prior to cytokine exposure (20).

### ROS, NO, and iron measurements

ROS levels were measured by DCF fluorescence (Abcam) as described (21). Nitrite levels were detected in supernatant culture media and measured using Griess reagents as described (22). NO/ROS measurements were normalized to total protein content (MicroBCA; ThermoFisher). Cellular labile iron pools were detected as described (23), imaged by confocal microscopy after loading with 1 µM calcein-AM (Invitrogen) and again following chelation with 300 µM deferasirox (Sigma).

### Western blot, reverse transcription, quantitative PCR, cell fractionation, and immunostaining

All assays were performed as previously described (11, 13). Commercially available antibodies used for Western blotting and immunostaining are listed in Supplementary Table 2. Rabbit Clec16a-specific polyclonal antisera (Cocalico) were generated following inoculation with a recombinant Clec16a protein fragment (AA347-472). Primer sequences used for quantitative reverse transcription PCR (qRT-PCR) are provided in Supplementary Table 3.

### Flow Cytometry

Following exposures and treatments, live mtKeima mouse islets were dissociated into single cells by the use of 0.25% trypsin-EDTA (Gibco) for 1min at 37°C. Single cells were stained with DAPI (Fisher) and re-suspended in phenol red-free RPMI1640 medium supplemented with 100 units/mL antimycotic-antibiotic (ThermoFisher), 50 units/mL Penicillin-Streptomycin (ThermoFisher), 1 mM sodium pyruvate (ThermoFisher) and 10 mM HEPES (ThermoFisher). Samples were analyzed on an LSR Fortessa flow cytometer (BD). Single cells were gated using FSC and SSC plots, and DAPI staining was used to exclude dead cells. Dual-excitation ratiometric mtKeima measurements were made using 488nm (neutral) and 561nm (acidic) excitation lasers with 610nm emission filter (24, 25). Ratios of acidic to neutral cell populations were then calculated using FlowJo (Treestar). 5,000 islet cells were quantified from each independent islet preparation.

Live Min6 β-cells transfected with tandem mitochondrial-targeted mCherry-eGFP mitophagy reporter (26) were detached from culture plates with the use of 0.25% trypsin-EDTA for 1min, re-suspended into single cells by gentle pipetting in phenol red-free DMEM media (Gibco) supplemented with 50 units/mL Penicillin-Streptomycin (ThermoFisher), 1 mM sodium pyruvate (ThermoFisher), and 142 µM β-mercaptoethanol, and then stained with DAPI. Samples were then analyzed on a ZE5 flow cytometer (BioRad). Single cells were gated using FSC and SSC plots, and DAPI staining was used to exclude dead cells. Mitochondrial fluorescence in cell populations was assessed using 488nm (GFP) and 561nm (mCherry) excitation lasers with 525nm and 610nm emission filter sets, respectively. Live cell populations were then gated for mCherry-positive cells that contained low GFP fluorescence (FlowJo) to quantify mitophagy. 10,000 cells were quantified from each independent sample.

### Mitochondrial bioenergetic measurements

Oxygen consumption was measured using oxygen micro-sensing glass electrodes (UniSense). The membrane potential (V_m_) of human islets was measured with patch electrodes having <10 Mohm resistance connected to micro-manipulators and a HEKA USB EPC10 patch-clamp amplifier (Heka Instruments) in the perforated patch-clamp configuration. Islets were perifused with a standard external solution containing 135 mM NaCl, 4.8 mM KCl, 3 mM CaCl_2_, 1.2 mM MgCl_2_, 20mM HEPES, 10 mM glucose; pH 7.35), or when higher KCl solutions were used, NaCl was reduced equimolar for KCl. β-cells were identified by the presence of characteristic slow oscillations in 10 mM glucose (27). ATP:ADP ratio was monitored using the Perceval-HR biosensor (28). Islets were perifused with 16.7 mM glucose to stimulate ATP production in a recording chamber on an Olympus IX-73 inverted epifluorescence microscope (Olympus). Perceval-HR was excited at 488nm using a TILL Polychrome V mono-chromator (FEI), and a QuantEM:512SC cooled CCD camera (PhotoMetrics) was used to collect emission at 527nm (20). Data were acquired and analyzed using Metafluor (Invitrogen) and plotted using Igor Pro (WaveMetrics Inc.).

Live-cell imaging in other studies was performed using a laser-scanning Nikon A1 confocal microscope equipped with an environmental chamber (Tokai Hit) maintained at 37°C and 5% CO_2_. Z-stack images were acquired using a Plan Fluor 40x oil objective (Nikon Instruments) and NIS Elements Software (Nikon Instruments). ΔΨ_m_ was assessed in live human islets using the membrane potential-sensitive pentamethinium fluorescent probe, TBMS-306 (29), a gift from Vladimir Kral and Tomas Ruml. Islets were stained with TBMS-306 (50nM) for 30min at 37 °C and co-stained with Fluozin-3 (500nM) to identify β-cells, as described (30). Z-stack images were acquired using 488nm (Fluozin-3) and 640nm (TBMS-306) excitation lasers with 510nm and 670nm emission filter sets, respectively.

### Cell death assessments

Cytoplasmic histone-complexed DNA fragments were determined in islets using the Cell Death ELISAplus (Roche), per the manufacturer’s protocol. Cell pellets were analyzed to detect apoptosis. ELISA absorbance was measured at 450nm and normalized to total DNA content (SYBRGreen; Invitrogen) using an absorbance/fluorescence microplate reader (BioTek). Terminal deoxy-nucleotidyl transferase dUTP nick end labeling (TUNEL) was performed on transduced human islets (Apoptag In situ Apoptosis Detection Kit; Millipore) as per the manufacturer’s protocol. Human islets were prepared for staining as previously described (11). TUNEL stained samples were counter-stained with insulin and GFP antibodies. Images were captured with an IX81 microscope (Olympus) using an ORCA Flash4 CMOS digital camera (Hamamatsu). Apoptotic transduced β-cells (TUNEL+, insulin+, GFP+) were counted as a fraction of total transduced β-cells (insulin+, GFP+). DAPI served as a nuclear DNA control.

### Mitochondrial morphology and sub-cellular localization

Mitochondrial morphology and localization studies were performed on immunostained human islets or paraffin-embedded mouse pancreas tissue sections, prepared as previously described (11, 13). Z-stack images were captured with an IX81 microscope (Olympus) using an ORCA Flash4 CMOS digital camera (Hamamatsu) and subjected to deconvolution (CellSens; Olympus). Co-localization analyses were performed using the Coloc2 plugin on ImageJ. Mitochondrial morphology was visualized using 3D-renderings generated with Imaris® imaging software (Bitplane). Quantitative 3D assessments of mitochondrial morphology and network were performed using the MitoMap and MitoAnalyzer plugins on ImageJ (31, 32).

### Statistics

Data are presented as means, and error bars denote SEM. Statistical comparisons were performed using unpaired Student’s t-test or one-way ANOVA as appropriate (Prism GraphPad). A *P* value < 0.05 was considered significant.

## Results

### Pro-inflammatory cytokines induce mitochondrial damage and activate β-cell mitophagy

Optimal mitochondrial function is central to β-cell responses to glucose or other nutrient stimuli. We hypothesized that pro-inflammatory cytokines induce mitochondrial dysfunction, and β-cells then activate mitophagy to eliminate dysfunctional mitochondria. To this end, we first examined the effects of pro-inflammatory cytokines (combination of IL-1β, TNF-α, and IFN-γ) on mitochondrial function in primary human islets. Mitophagy is initiated following a loss of mitochondrial membrane potential (ΔΨ_m_) and resultant respiratory dysfunction (13, 33). Utilizing live-cell confocal microscopy, we observed that cytokine exposure dissipated ΔΨ_m_ primarily in β-cells, which were detected by the cell permeable Zn^2+^ dye Fluozin-3 (Figure 1A; (30)). Moreover, cytokine exposure reduced both oxygen consumption (Figure 1B) and ATP/ADP ratio (Figure 1C) of human islets in response to glucose stimulation. Glucose-induced increases in the ATP/ADP ratio are necessary for closure of ATP-sensitive potassium (K_ATP_) channels to produce β-cell membrane depolarization, and indeed, patch clamping confirmed that cytokine exposure reduced glucose-stimulated membrane depolarization (Figure 1D). However, β-cell depolarization was still seen in response to the sulfonylurea tolbutamide, which closes K_ATP_-channels independently of glucose metabolism, suggesting that the effects of cytokines are metabolic, and thus occur upstream of the K_ATP_-channel (Figure 1E). Together, these studies confirm that pro-inflammatory cytokines induce mitochondrial dysfunction in human β-cells.

**Figure 1.**
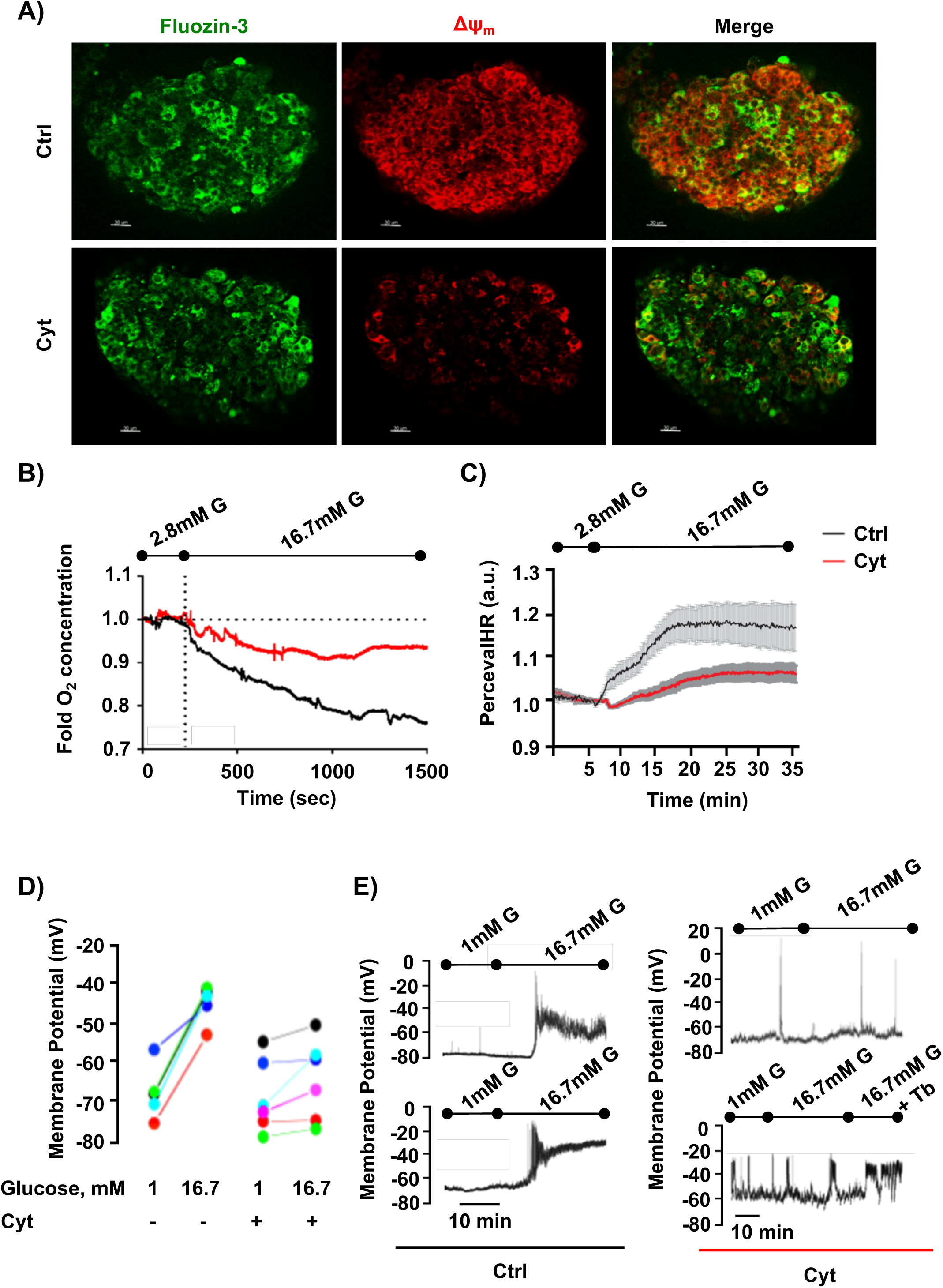
Pro-inflammatory cytokines impair mitochondrial bioenergetics and metabolic function in human islets. **(A)** Laser scanning confocal microscopy of live human islets stained with Fluozin-3 (β-cells/Zn granules) and TBMS-306 (ΔΨ_m_) following a 24-hr treatment with control (Ctrl; PBS) or cytokines (Cyt; 75 U/mL IL-1β, 750 U/mL TNF-α, and 750 U/mL IFN-γ). **(B)** O_2_ consumption measured by O_2_ microsensor in Ctrl and Cyt-treated human islets (p<0.05 by ANOVA). **(C)** ATP:ADP ratios measured by Perceval-HR fluorescence in Ctrl and Cyt-treated human islets (p<0.05 by ANOVA). **(D)** Membrane potential assessments by patch clamp in Ctrl and Cyt-treated human islets. Colors represent measurements from independent human islet donors. **(E)** Representative electrical recordings from Ctrl (left) and Cyt-treated (right) human islets stimulated with glucose and 100μM tolbutamide (Tb) as indicated. n=3-6 independent human islet donors/group for all measurements.

The initiation of mitophagy is marked by recruitment of the cytosolic E3 ligase Parkin to depolarized mitochondria, resulting in turnover of outer mitochondrial membrane (OMM) proteins including mitofusins 1 and 2 (Mfn1 and Mfn2), turnover of Parkin itself, and then clearance of damaged mitochondria by the autophagosome-lysosome pathway (33). In Min6 β-cells exposed to inflammatory cytokines, endogenous Parkin translocated to the mitochondria (Figure 2A).

**Figure 2.**
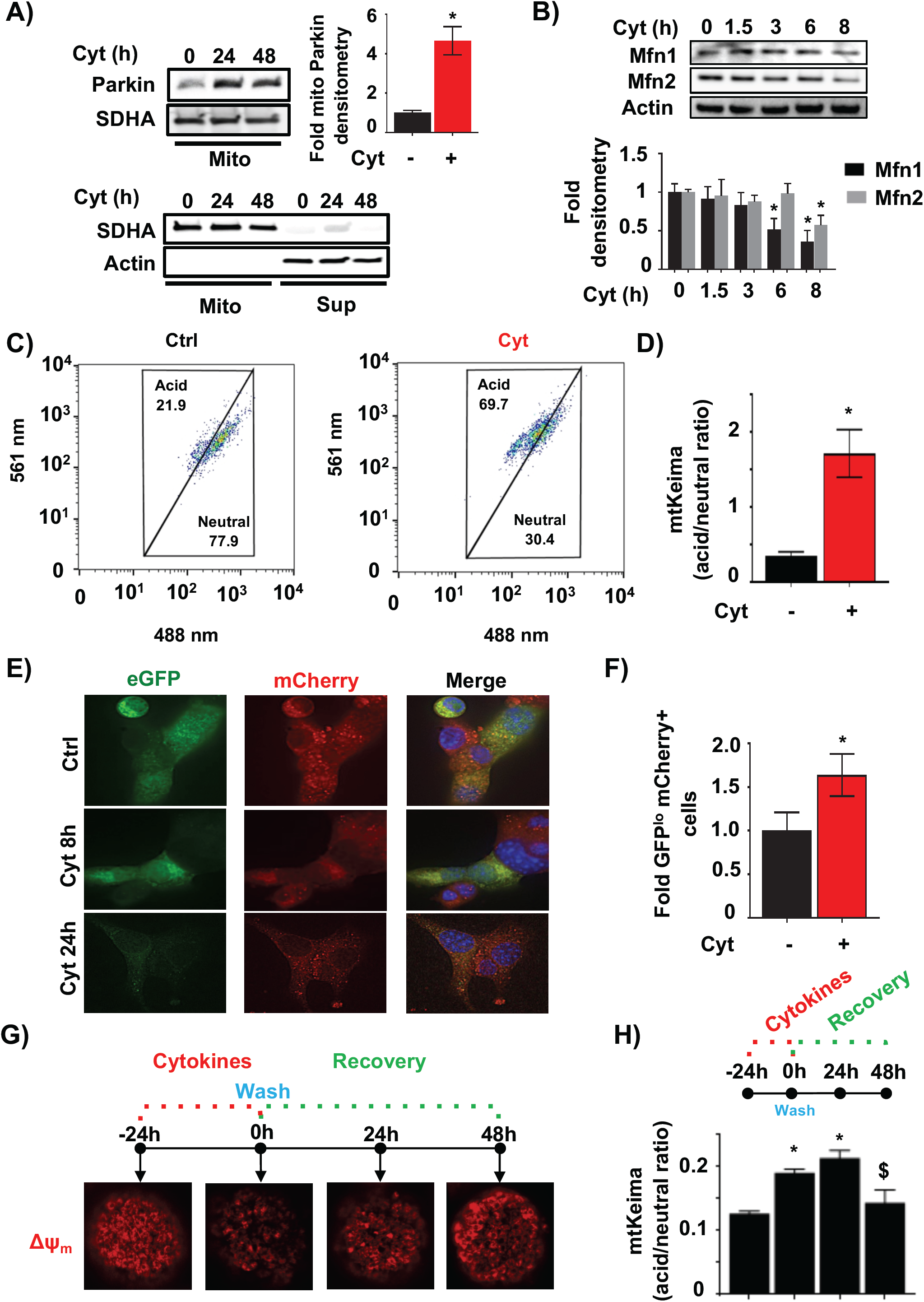
Mitophagy is activated by pro-inflammatory cytokines. **(A)** <u>Top</u>: Mitochondrial Parkin localization by Western blot (WB), with densitometry normalized to SDHA as a loading control in cytokine-treated Min6 β-cells following biochemical fractionation of mitochondria. <u>Bottom</u>: WB following cell fractionation and 8000g centrifugation of cytokine (or vehicle)-treated Min6 β-cells probed for SDHA (mitochondria; mito) or actin (supernatant; sup) to detect purity of mitochondrial fraction. * p<0.05. **(B)** Mfn1 and Mfn2 expression by WB (with densitometry normalized to actin) in Min6 β-cells treated with cytokines for indicated time course. * p<0.05 vs. Cyt 0hr. **(C)** Flow cytometry scatter plot from FVB/N mtKeima islets treated with control or cytokines for 6hrs. Cells gated in upper acid group represent populations with increased relative acidic 561nm to neutral 480nm excitation, consistent with activation of mitophagy. **(D)** Quantification of flow cytometry data by acid/neutral population ratio from FVB/N mtKeima islets in Figure 2C treated with control or cytokines for 6hrs. * p<0.05. **(E)** Deconvolution image of live Min6 β-cells transfected with mitochondria-targeted tandem mCherry-eGFP plasmid following control or cytokine exposure for indicated time course. Nuclei are stained by Hoechst 33342 (blue). **(F)** Flow cytometric quantification of eGFP and mCherry fluorescence of live Min6 β-cells transfected as in Figure 2E following control or cytokine treatment for 24hrs. Data expressed as fold change of eGFP low and mCherry positive cell populations to indicate mitophagy. * p<0.05. **(G)** Laser scanning confocal microscopy of live B6N mouse islets stained with TBMS-306 (ΔΨ_m_) following cytokine treatment for 24hrs and recovery after cytokine removal for 24 and 48hrs. **(H)** Flow cytometric quantification of acid/neutral population ratio from B6N mtKeima islets following cytokine treatment for 24hrs and recovery after cytokine removal for 24 and 48hrs. * p<0.05 vs. - 24hr-control islets. $ p<0.05 vs. 0hr-islets. n=3-6/group for all studies.

Furthermore, we observed a time-dependent decrease of Mfn1 and Mfn2 protein following cytokine exposure (Figure 2B). Classical inducers of mitophagy, including FCCP and valinomycin, induced similar turnover of Mfn1 and Mfn2 protein (Figure S1A). Importantly, cytokines induced β-cell mitophagy but not bulk macroautophagy, as we neither observed differences in the protein levels or cleavage/activation of LC3 (Figure S1B), nor in the protein levels of the autophagy substrate p62 following cytokine exposure in mouse islets (data not shown).

To further assess mitophagy following cytokine exposure, we utilized fluorescently labeled and pH-sensitive mitochondrial biosensors to analyze the translocation of mitochondria to acidic lysosomes via two independent and previously validated approaches (17, 26). We first analyzed mitophagy rates in primary islets isolated from mtKeima mice which express a mitochondria-targeted reporter that exhibits a shift in excitation/emission spectra based on changes in pH (17). Notably, mtKeima mice do not exhibit any defects in glucose tolerance when compared to littermate transgene-negative controls (data not shown). Flow cytometry of dissociated islets revealed that cytokine exposure increased the number of cells whose mitochondria were localized to acidic compartments (Figures 2C-D). As a complementary approach, we expressed a mitochondria-targeted tandem mCherry-eGFP reporter in Min6 β-cells (Figure 2E). In cells expressing this reporter, both eGFP and mCherry are detectable in mitochondria at neutral pH, but when mitochondria are in acidic lysosomes, the eGFP fluorescence is quenched (26). Following cytokine exposure, we observed a reduction in eGFP but not mCherry fluorescence, consistent with localization of damaged mitochondria to lysosomes (Figures 2E-F).

As β-cells can recover from inflammatory damage following the withdrawal of cytokines (34), we asked if ΔΨ_m_ and mitophagy would return to baseline following removal of inflammatory stress. Indeed, cytokine withdrawal rescued ΔΨ_m_ in WT mouse islets, and normalized rates of mitophagy in mtKeima mouse islets (Figures 2G-H). Taken together, these studies indicate that β-cells activate mitophagy in response to inflammatory stress.

### NO and/or ROS are inflammatory mediators of cytokine-induced mitophagy

Pro-inflammatory cytokines activate signaling cascades that have both protective and detrimental outcomes. Among the most well-known effectors of inflammatory cytokines are nitric oxide (NO) and reactive oxygen species (ROS) (6). To determine the role of free radicals in the induction of mitophagy, we treated mtKeima islets and tandem mito-mCherry-GFP expressing Min6 β-cells with pharmacological agents to increase NO or ROS, including DPTA/NO (a NO donor), paraquat (which induces mitochondrial superoxide production), or rotenone (a mitochondrial complex I inhibitor which induces ROS formation). In both islets and Min6 β-cells, induction of NO or ROS activated mitophagy (Figure 3A) to a similar extent as valinomycin, a potassium ionophore that induces mitophagy by dissipating ΔΨ_m_. Further, we observed a concordant decrease in OMM proteins and total Parkin levels upon exposure to rotenone and/or DPTA/NO, again consistent with the induction of β-cell mitophagy by ROS or NO (Figures 3B-D). To determine if inflammatory cytokines induce mitophagy via free radical generation, we utilized both genetic and pharmacologic approaches. β-cells generate cytokine-induced NO via inducible nitric oxide synthase (iNOS, encoded by *NOS2*; (35, 36)); therefore, we intercrossed *NOS2*^-/-^ mice and mtKeima mice for mitophagy analysis. As expected, loss of iNOS decreased NO release following cytokine exposure in islets (Figures 3E-F). In addition, cytokine-mediated induction of mitophagy was abrogated in the islets of *NOS2*^-/-^;mtKeima mice (Figure 3G). Similarly, treatment with the mitochondria-permeable superoxide scavenger, tiron (37) prevented activation of cytokine-induced mitophagy in wild-type mtKeima islets (Figure 3G). Of note, tiron may have effects to scavenge NO in addition to superoxide (38, 39). Thus, our results indicate that cytokine-induced free radicals elicit mitophagy in β-cells.

**Figure 3.**
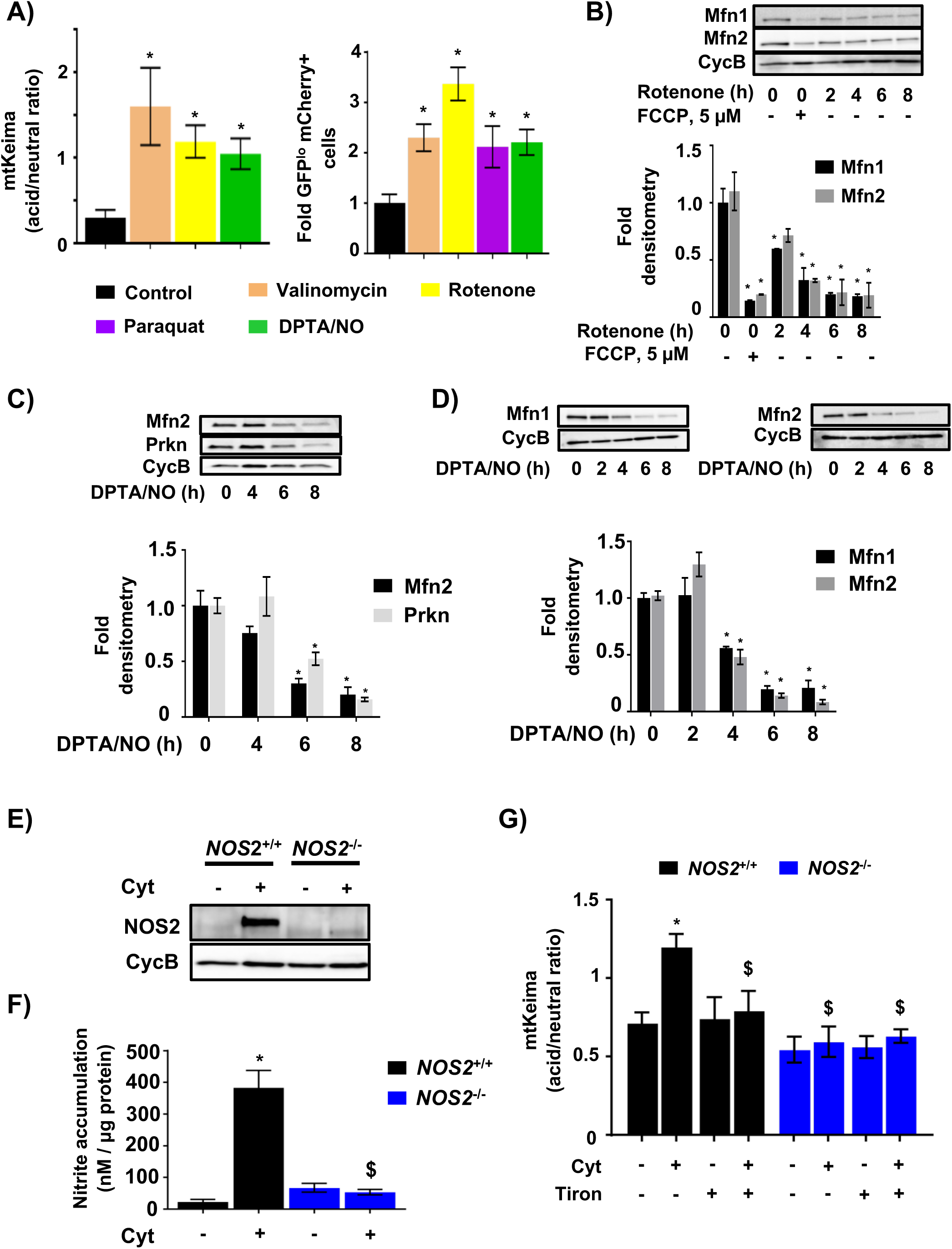
Nitric oxide/reactive oxygen species regulate cytokine-mediated mitophagy. **(A)** <u>Left</u>: Assessment of mitophagy by flow cytometric quantification of acid/neutral ratio from FVB/N mtKeima islets following treatment with Valinomycin (250nM for 3hrs), Rotenone (250nM for 3hrs) and DPTA/NO (600µM for 1hr). * p<0.05 vs. control islets. <u>Right</u>: Flow cytometric quantification of eGFP and mCherry fluorescence of Min6 β-cells transfected with mitochondria-targeted tandem mCherry-eGFP mitophagy reporter following treatment with Valinomycin (250nM for 4hrs), Rotenone (500nM for 4hrs), Paraquat (1mM for 4hrs) and DPTA/NO (600µM, 6hrs). n=3/group; * p<0.05 vs. control cells. **(B)** Mfn1 and Mfn2 expression by WB (with densitometry normalized to cyclophilin B) in Min6 β-cells treated with 5 µM FCCP for 6hrs or 500nM Rotenone for indicated time course. * p<0.01 vs. 0hr-control. **(C)** Mfn2 and Prkn expression by WB (with densitometry normalized to cyclophilin B) in mouse islets treated with DPTA/NO (600µM) for indicated time course. * p<0.05 vs. 0hr-control. **(D)** Mfn1 and Mfn2 expression by WB (with densitometry normalized to cyclophilin B) in Min6 β-cells treated DPTA/NO (600µM) for indicated time course. * p<0.05 vs. 0hr-control. **(E)** NOS2 expression by WB in *NOS2*^+/+^ and *NOS2*^-/-^ islets treated with Ctrl (PBS) or Cyt for 24hrs. **(F)** Secreted nitrite concentrations (normalized to islet total protein content) in supernatant collected from cultured *NOS2*^+/+^ and *NOS2*^-/-^ islets treated with PBS or Cyt for 24hrs. * p<0.05 vs. PBS-treated WT islets. $ p<0.05 vs. Cyt-treated WT islets. **(G)** Assessment of mitophagy by flow cytometric quantification of acid/neutral ratio from *NOS2*^+/+^;mtKeima (Black) and *NOS2*^-/-^;mtKeima (Blue) islets treated with 500µM tiron for 24hrs with cytokines for the final 6hrs. * p<0.05 vs. WT PBS-treated islets. $ p<0.05 vs. WT Cyt-treated islets. n=3-5/group for all studies.

### Impaired β-cell mitophagy exacerbates hyperglycemia and mitochondrial fragmentation *in vivo* following inflammatory stimuli

Our observation that mitophagy is activated following inflammatory β-cell damage led us to hypothesize that mitophagy is a protective response that preserves β-cell function in the setting of inflammatory stress. To interrogate the role of mitophagy *in vivo*, we deleted the key mitophagy regulator Clec16a specifically in β-cells (*Clec16a*^loxP/loxP^; *Ins1*-Cre, hereafter referred to as β-Clec16a^KO^; Figure 4A). *Clec16a* encodes an E3 ubiquitin ligase vital for the clearance of dysfunctional β-cell mitochondria via mitophagy (11, 12). As expected, unchallenged β-Clec16a^KO^ mice did not exhibit defects in random fed blood glucose values and only mild glucose intolerance after an intraperitoneal glucose challenge (data not shown), consistent with our previous reports (11, 13).

**Figure 4.**
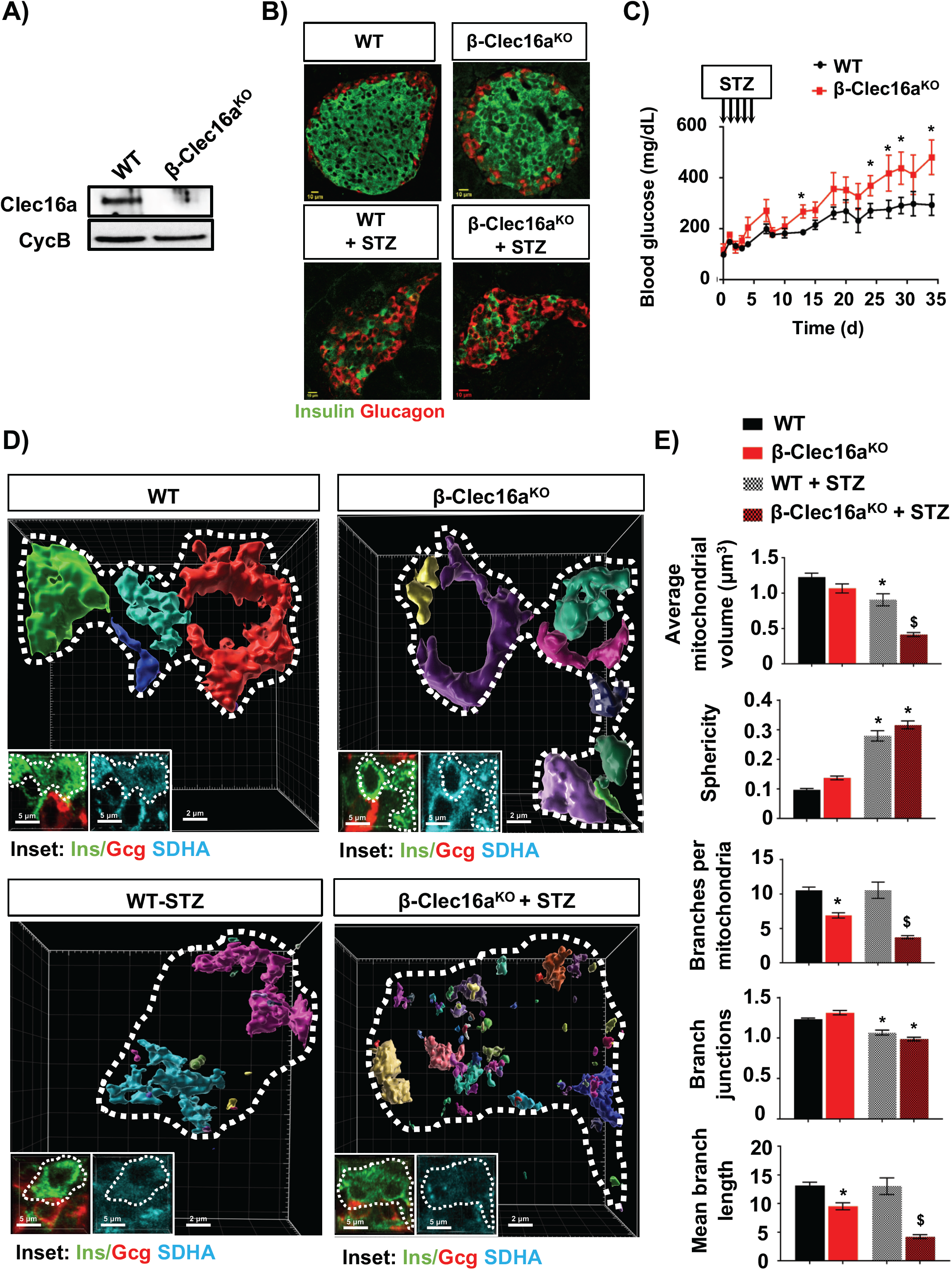
Mitophagy deficiency aggravates STZ-induced dysglycemia and mitochondrial network defects *in vivo*. **(A)** Clec16a expression by WB in 10-week-old WT and β-Clec16a^KO^ mouse islets. Cyclophilin B serves as a loading control. **(B)** Random fed blood glucose concentrations from WT (Black) and β-Clec16a^KO^ mice (Red) following STZ treatment. *p<0.05 vs. WT. **(C)** Immunofluorescence images of insulin (green) and glucagon (red) from pancreatic sections of WT and β-Clec16a^KO^ mice, in the presence or absence of STZ treatment (50 mg/kg/d IP for 5d). **(D)** Imaris generated three-dimensional reconstruction of confocal immunofluorescence Z-stack images stained for SDHA (see inset image – blue) from pancreatic sections of WT and β-Clec16a^KO^ mice, in the presence or absence of STZ treatment. β-cells and α-cell were identified by co-staining (inset: insulin – Ins;green, glucagon – Gcg; red). β-cells are encircled by a dotted line (white). Each unique color represents a separate β-cell mitochondrial network cluster. **(E)** β-cell mitochondrial morphology and network analysis of confocal immunofluorescence Z-stack images from studies depicted in Figure 4D, stained for SDHA (and insulin) from pancreatic sections of WT and β-Clec16a^KO^ mice, in the presence or absence of STZ treatment by MitoAnalyzer. (185-220 β-cells/animal were quantified). *p<0.05 vs. WT. $ p<0.05 vs. WT + STZ. n=4-8 mice/group for all studies.

To determine the importance of mitophagy in β-cell inflammatory responses, we treated β-Clec16a^KO^ mice and littermate controls with multiple low doses (50 mg/kg/d x 5 days) of the β-cell toxin and inflammatory stressor streptozotocin (STZ) (40-42). Both WT and β-Clec16a^KO^ mice showed a reduction in β-cells in response to STZ (Figures 4B). However, the effect of STZ on blood glucose levels was significantly exacerbated in β-Clec16a^KO^ mice (Figure 4C), supporting a role for mitophagy in preserving β-cell function following inflammatory stress.

Damaged mitochondria are segregated from the mitochondrial network by fission, leading to a fragmented appearance of the network prior to the elimination of dysfunctional mitochondria by mitophagy (43). We hypothesized that inflammatory stress due to STZ would induce mitochondrial fragmentation, which would be exacerbated in mitophagy-deficient β-cells. We thus examined mitochondrial morphology and networks in islets of β-Clec16a^KO^ mice and littermate controls in the presence or absence of STZ. Utilizing confocal imaging of the mitochondrial marker succinate dehydrogenase A (SDHA), we generated three-dimensional (3D) reconstructions of islet mitochondria and quantified mitochondrial morphology and network appearance. In the absence of STZ, mitochondria from control and β-Clec16a^KO^ β-cells revealed relatively similar 3D mitochondrial appearance between genotypes (Figure 4D), with a mild reduction in branch complexity in β-Clec16a^KO^ mitochondria compared to controls (Figure 4E). STZ-treatment, in contrast, increased the frequency of smaller vesiculated mitochondrial networks in WT β-cells consistent with fragmentation; however, mitochondrial fragmentation was dramatically exacerbated by the deletion of Clec16a (Figure 4D). 3D quantification confirmed that STZ-treatment altered mitochondrial morphology and network complexity in control β-cells, decreasing mitochondrial volume and branch junctions and increasing their sphericity (Figure 4E). Mitochondrial morphology and network/branch complexity markedly worsened in STZ-treated β-Clec16a^KO^ β-cells, consistent with an accumulation of fragmented mitochondria due to deficient mitophagy. Together, these data support an important role for mitophagy in protecting β-cell mitochondrial networks from inflammatory stimuli *in vivo*.

### Clec16a regulates cytokine-induced mitophagy in rodent and human islets

The accumulation of fragmented mitochondria in β-cells from STZ-treated Clec16a-deficient mice led us to speculate that Clec16a regulates mitochondrial turnover following exposure to pro-inflammatory cytokines. Indeed, β-Clec16a^KO^ islets had impaired clearance of Mfn2 and Parkin following cytokine exposure, consistent with Clec16a control of cytokine-induced mitophagy (Figures 5A-B). Similarly, treatment of islets from mtKeima mice with lenalidomide, a pharmacologic inhibitor of the CLEC16A-mediated mitophagy pathway (11, 44), impaired cytokine-induced mitophagy (Figure 5C).

**Figure 5.**
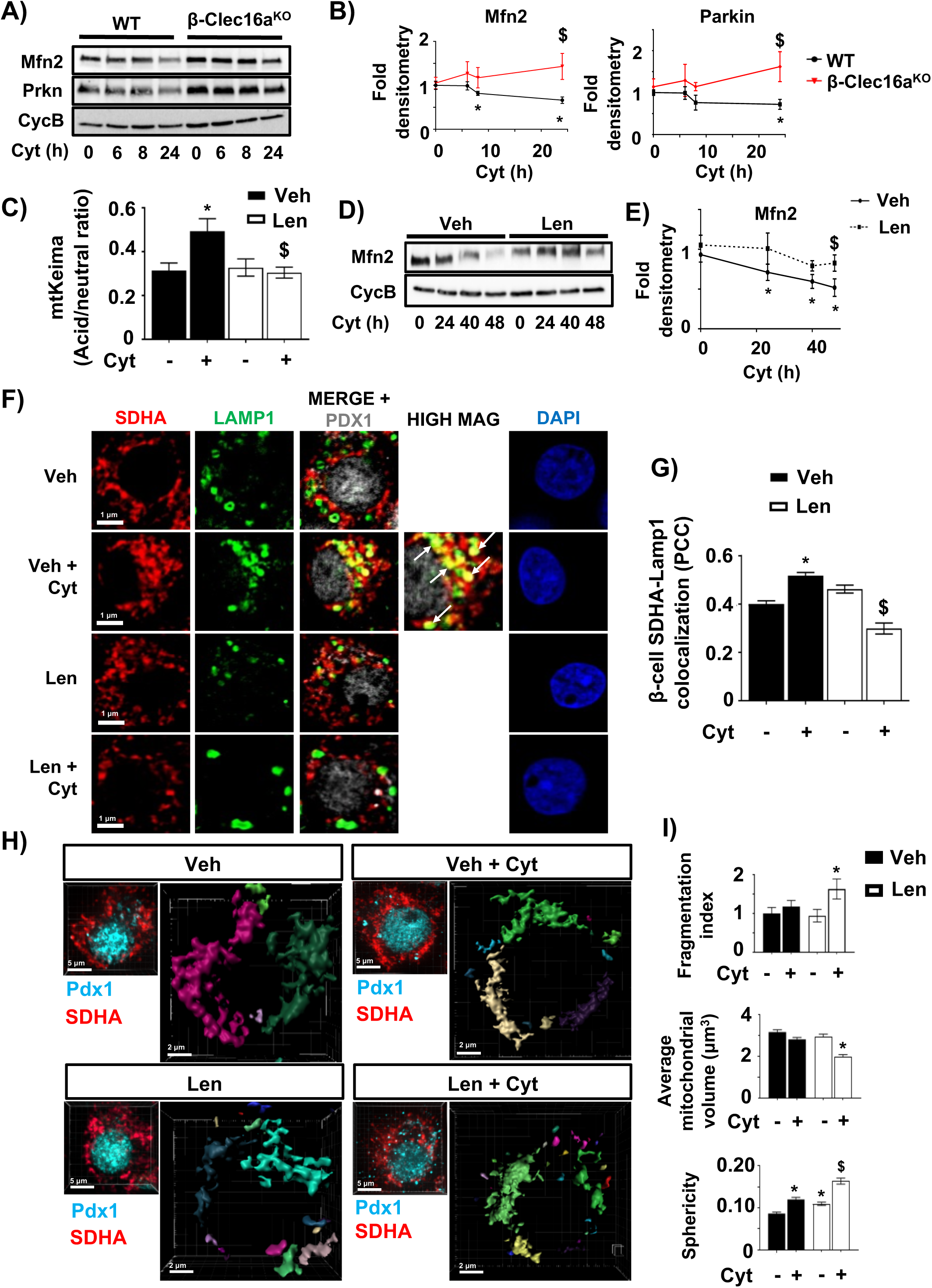
Clec16a regulates cytokine-induced mitophagy in human and rodent islets. **(A)** Mfn2 and Prkn expression by WB in WT and β-Clec16a^KO^ islets treated with cytokines for indicated time course. **(B)** Mfn2 and Prkn densitometry (normalized to Cyclophilin B) from studies in Figure 5A. * p<0.05 vs. WT 0hr; $ p<0.05 vs. WT 24hrs. **(C)** Assessment of mitophagy by flow cytometric quantification of acid/neutral ratio from B6N mtKeima islets treated with vehicle (Veh; DMSO) or 10µM lenalidomide (Len) for 24hrs with cytokines for the final 6hrs. * p<0.05 vs. Veh alone; $ p<0.05 vs. Veh + Cyt. **(D)** Mfn2 expression by WB in human islets treated with Veh or 10µM Len for 72hrs in the presence of cytokines for the final 48hrs per indicated time course. **(E)** Mfn2 densitometry (normalized to Cyclophilin B) from studies in Figure 5D. * p<0.05 vs. Veh 0hr; $ p<0.05 vs. Veh 48hrs. **(F)** Confocal immunofluorescence image of human islets treated with Veh or 10µM Len for 72hrs in the presence/absence of cytokines for the final 24hrs stained for SDHA (red), LAMP1 (green), Pdx1 (gray), and DAPI (blue). **(G)** Quantification of Lamp1^+^SDHA^+^ colocalization in human β-cells from studies depicted in Figure 5F by Pearson’s correlation coefficient (PCC). * p<0.05 vs. Veh + PBS; $ p<0.05 vs. Veh + Cyt. (60-130 β-cells from each donor per condition were analyzed) **(H)** Imaris generated three-dimensional renderings of confocal immunofluorescence Z-stack images stained for SDHA (see offset images – red) in human β-cells from studies depicted in Figure 5F. β-cells were identified by Pdx1 co-staining (see offset images – blue). Each unique color represents a separate β-cell mitochondrial network cluster. **(I)** Human β-cell mitochondrial morphology and network analysis of confocal immunofluorescence Z-stack images stained for SDHA from studies depicted in Figure 5F by Mitomap and MitoAnalyzer. (75-110 β-cells from each donor per condition were quantified). *p<0.05 vs. Veh + PBS. $ p<0.05 vs. Veh + Cyt. n=3-5/group for all studies.

To interrogate mitophagy as a response to inflammatory damage in human islets, we assessed several complementary mitochondrial endpoints. Cytokine exposure led to reduced Mfn2 protein levels as well as increased localization of mitochondria (marked by SDHA) within lysosomes (marked by LAMP1), consistent with the activation of mitophagy (Figures 5D-G). Cytokine-induced mitophagy was also blocked by lenalidomide treatment (Figures 5D-G). Furthermore, the combination of cytokines and lenalidomide increased mitochondrial fragmentation (Figures 5H-I and Figure S2). Therefore, in both human and rodent β-cells, CLEC16A-mediated mitophagy protects mitochondrial network integrity following inflammatory stress.

### Mitophagy deficiency exacerbates cytokine-induced β-cell death

While we demonstrated that mitophagy preserved mitochondrial networks following inflammatory injury, it was unclear if mitophagy also preserved β-cell viability after inflammatory damage. To address this question, we assessed apoptosis in Clec16a-deficient β-cells (and controls) following cytokine exposure. Indeed, cytokine exposure induced increased cell death in β-Clec16a^KO^ islets compared to controls (Figure 6A). Cytokine exposure similarly increased cell death in Min6 β-cells following shRNA-mediated Clec16a knockdown (Figures 6B-C and Figure S3A). Clec16a mRNA levels were unaffected by cytokine exposure in control samples (Figure S3A).

**Figure 6.**
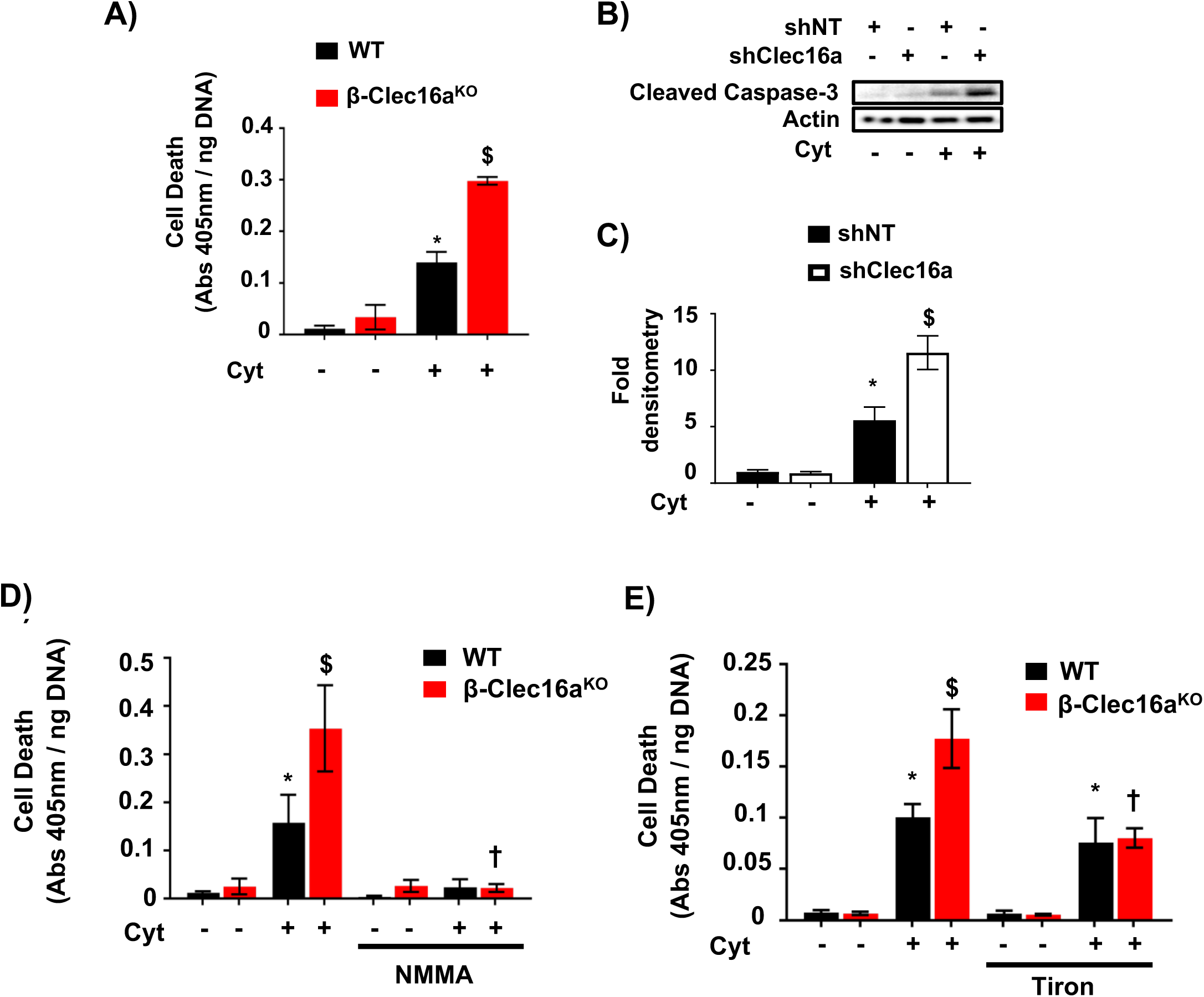
Mitophagy is a protective response to cytokine-induced nitrosative/oxidative stress. **(A)** Quantification of cell death by cytoplasmic histone-complexed DNA fragment ELISA (normalized to total DNA content) in WT and β-Clec16a^KO^ islets treated with/without cytokines for 24hrs. * p<0.05 vs. PBS-treated WT islets. $ p<0.05 vs. Cyt-treated WT islets. **(B)** Cleaved caspase 3 expression by WB in non-targeting (NT) or Clec16a-specific shRNA expressing Min6 β-cells treated with cytokines for 6hrs. **(C)** Cleaved caspase 3 densitometry (normalized to actin) from studies depicted in Figure 6B. * p<0.05 vs. shNT + PBS; $ p<0.05 vs. shNT + Cyt. **(D)** Quantification of cell death by cytoplasmic histone-complexed DNA fragment ELISA (normalized to total DNA content) in WT and β-Clec16a^KO^ islets treated with/without NMMA (500µM) for 48hrs and treated with/without cytokines for the final 24hrs. * p<0.05 vs. PBS-treated WT islets; $ p<0.05 vs. Cyt-treated WT islets; † p<0.05 vs. Cyt-treated β-Clec16a^KO^ islets. **(E)** Cell death ELISA (normalized to total DNA content) in WT and β-Clec16a^KO^ islets treated with/without 500µM tiron for 48hrs and treated with/without cytokines for the final 24hrs. * p<0.0005 vs. PBS-treated WT islets; $ p<0.05 vs. Cyt-treated WT islets; † p<0.05 vs. Cyt-treated β-Clec16a^KO^ islets. n=4-5/group for all studies.

Next, we utilized pharmacologic agents targeting NO/ROS to determine if free radicals contribute to the enhanced cytokine-induced apoptosis we observed in mitophagy-deficient islets. Indeed, treatment with the NOS inhibitor L-NMMA abrogated cytokine-induced apoptosis in both control and β-Clec16a^KO^ islets (Figure 6D). We also observed that enhanced cell death in β-Clec16a^KO^ islets was ameliorated by tiron (Figure 6E). Thus, mitophagy promotes β-cell survival in response to nitrosative/oxidative stress.

We next asked whether mitophagy-deficient β-cells are sensitized to cytokine-mediated apoptosis due to increased formation of free radicals. Interestingly, loss of Clec16a led to reduced, not increased, cytokine-induced nitrite levels compared to controls (Figure S3B). Further, Clec16a loss of function did not affect baseline or cytokine-induced ROS levels or expression of transcriptional targets associated with inflammatory NFκB signaling (Figures S3C-D). Moreover, Clec16a-deficiency did not affect basal or cytokine-induced cellular import of iron, a key catalyst of ROS production (Figure S3E). Similarly, inhibition of CLEC16A with lenalidomide did not affect cytokine-induced expression of NFκB targets in human islets (Figures S3F-G). Thus, we speculate that mitophagy mediates β-cell mitochondrial responses to nitrosative/oxidative stress, but does not directly affect the signaling pathways leading to free radical formation.

### CLEC16A overexpression protects human β-cells from cytokine toxicity

Our observations of enhanced cytokine-mediated apoptosis in mitophagy-deficient mouse islets led us to hypothesize that CLEC16A protects against β-cell death in human islets. Indeed, we observed that CLEC16A inhibition by lenalidomide enhanced cytokine-induced cell death in human islets (Figures 7A-B). In addition, NO/ROS blockade specifically ameliorated lenalidomide-related cell death, but did not affect vehicle-treated human islets (Figures 7A-B). To test whether CLEC16A overexpression could protect against human β-cell apoptosis, we transduced intact human islets with adenoviruses overexpressing CLEC16A or an empty vector control, as well as IRES-eGFP, selectively in β-cells (Figures S4A-B). Remarkably, CLEC16A overexpression significantly prevented cytokine-mediated apoptosis in human β-cells, even in the setting of higher IL-1β concentrations (Figures 7C and S4C). We also observed a similar protective effect of Clec16a overexpression in Min6 β-cells (Figures S4D-E). These results indicate that the mitophagy regulator CLEC16A protects human β-cells against inflammatory injury.

**Figure 7.**
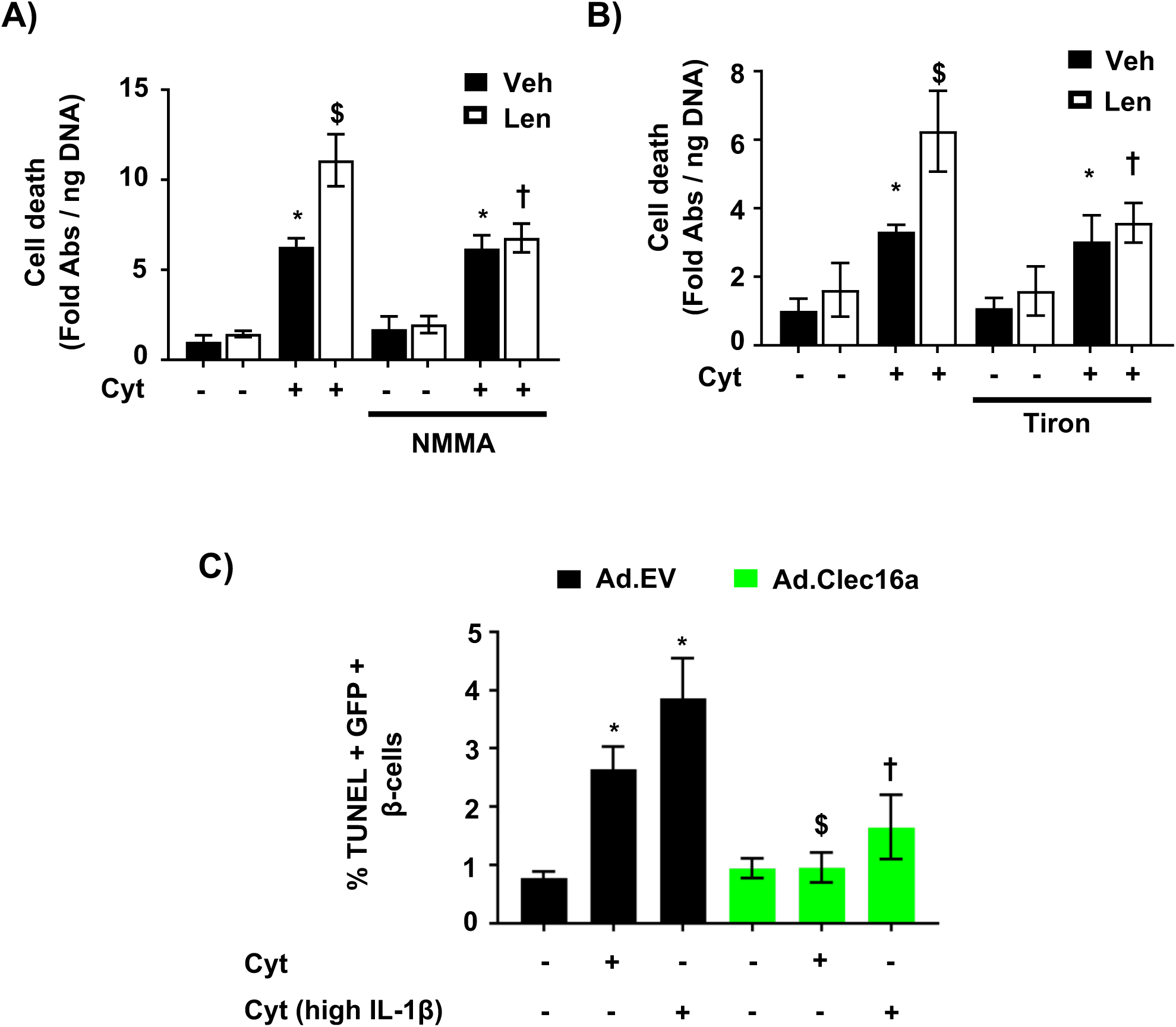
CLEC16A protects human β-cells from cytokine-induced apoptosis. **(A)** Cell death ELISA in human islets treated with/without 10µM lenalidomide and/or 500µM NMMA for 48hrs and then treated with/without cytokines for the final 24hrs. * p<0.05 vs. Veh + PBS; $ < 0.05 vs. Veh + Cyt; † p<0.05 vs. Len + Cyt. **(B)** Cell death ELISA in human islets treated with/without 10µM lenalidomide and/or 500µM tiron for 48hrs and then treated with/without cytokines for the final 24hrs. * p<0.05 vs. Veh + PBS; $ p<0.05 vs. Veh + Cyt; † p<0.05 vs Len + Cyt. **(C)** Quantification of %TUNEL^+^ GFP^+^ β-cells by immunofluorescence staining on human islets transduced with empty vector control (Ad.EV; black) or Clec16a-overexpressing (Ad.Clec16a; green) adenoviral particles, following treatment with/without cytokines for 24hrs. β-cells were identified by insulin immunostaining, and transduced cells were identified by GFP immunostaining. Cyt group treated with 75U/mL IL-1β, 750 U/mL TNFα, and 750 U/mL IFNγ. Cyt-high IL-1β group treated with 3,000 U/mL IL-1β, 750 U/mL TNFα, and 750 U/mL IFNγ. * p<0.05 vs. Ad.EV + PBS, $ p<0.05 vs. Ad.EV + Cyt, † p<0.05 vs. Ad.EV + Cyt high IL-1β. n=4-5/group for all studies.

## Discussion

Here we identify mitophagy as a vital protective response to inflammatory assault of human β-cells. We demonstrate that inflammatory cytokines activate the cardinal steps of mitophagy, including the dissipation of ΔΨ_m_, mitochondrial translocation of Parkin, turnover of OMM proteins, mitochondrial segregation, and mitochondrial localization to lysosomes for their elimination. We show that cytokine-mediated mitophagy is activated in response to free radicals. Moreover, we identified a protective role for the T1D gene *CLEC16A* in preventing β-cell death following inflammatory stress.

Our results indicate that by controlling mitophagy, CLEC16A may be a key determinant of β-cell susceptibility to inflammation-induced apoptosis. Recent reports position CLEC16A as the principal mediator of T1D risk at the chromosome 16.13 locus (13, 45-47). While *CLEC16A* polymorphisms in T1D are associated with reduced human islet CLEC16A expression and impaired β-cell function and glycemic control, prior studies have not clarified the importance of CLEC16A in β-cells following diabetogenic stressors. Importantly, we observe that CLEC16A-deficiency sensitizes β-cells to inflammatory stress, which may provide insight into how *CLEC16A* polymorphisms increase susceptibility to the development of T1D. Future studies in autoimmune diabetes models will be necessary to further clarify β-cell specific roles for CLEC16A in T1D. However, our findings lead us to speculate that CLEC16A deficiency may be a novel mediator of β-cell fragility (48, 49), which has been recently appreciated as a component of T1D pathogenesis.

*CLEC16A* is ubiquitously expressed and also has key functions in immune cells (50-55). Recent work demonstrating mitochondrial metabolic defects linked to inflammatory responses in immune cells of T1D donors may suggest an unexplored role for CLEC16A-mediated mitophagy in T1D (7, 56). Likewise, defective mitophagy has been associated with elevated secretion of cytokines by macrophages (57, 58). Together with our studies in β-cells, CLEC16A-mediated mitophagy may play pleotropic roles in multiple cell types in T1D and therefore may represent a potent therapeutic target.

Our studies are the first to show a specific and selective link between mitophagy and protection against inflammatory β-cell death. Autophagy comprises several pathways that maintain cellular homeostasis, including selective and non-selective/bulk macroautophagy, as well as microautophagy and chaperone-mediated autophagy (59). Previous studies on macroautophagy demonstrate a cytoprotective role in β-cells (60, 61), but these studies primarily relied upon β-cell specific deletion of proteins that broadly impair both selective and non-selective autophagy. Therefore, the importance of non-selective versus selective autophagy in β-cells has remained unresolved. Non-selective macroautophagy is classically induced by nutrient deprivation (62), whereas diabetes is a disease of glucose/nutrient excess, suggesting an underappreciated role for other forms of autophagy in β-cell function and survival. Importantly, our data demonstrate a specific induction of mitophagy (and not bulk macroautophagy) by inflammatory damage, potentially placing selective autophagy in a key position in the development of diabetes.

Our studies demonstrate a protective role of CLEC16A-mediated mitophagy in human β-cells exposed to inflammatory stimuli and support the feasibility of therapeutically targeting this pathway. Several mitophagy activating compounds have been recently found to improve respiratory function in metabolic tissues, including β-cells (63-66). Indeed, the mitophagy activator urolithin A has shown great promise in support of whole body metabolic function in clinical trials (67). Therefore, pharmacologic enhancement of mitophagy could represent a novel future approach to prospectively compensate for inflammatory stress and prevent β-cell death in diabetes. Future studies will be required to fully characterize the efficacy of targeting mitophagy in the treatment of all forms of diabetes.

## Acknowledgements

Human pancreatic islets were provided by the NIDDK-funded Integrated Islet Distribution Program (IIDP) at City of Hope, NIH Grant # 2UC4DK098085. S.A.S was supported by the JDRF (CDA-2016-189, SRA-2018-539, COE-2019-861), the NIH (R01 DK108921), the Department of Veterans Affairs (I01 BX004444), the Brehm family, and the Anthony family. G.L.P. was supported by the American Diabetes Association (19-PDF-063). V.S.P. was supported by an Upjohn postdoctoral fellowship. L.C. and T.M-P. were supported by the JDRF (CDA-2016-189). M.A.G. was supported by the NIH (T32-GM008322 and F31-DK122761). L.S.S. was supported by the NIH (R01 DK46409) and the JDRF (SRA-2018-539). The JDRF Career Development Award to S.A.S. is partly supported by the Danish Diabetes Academy and the Novo Nordisk Foundation. We acknowledge Dr. S. Lentz of the Microscopy, Imaging and Cellular Physiology Core of the University of Michigan DRC (P30DK020572) for assistance with imaging studies. We thank the University of Michigan Flow Cytometry Core for assistance with flow cytometry studies. We thank Drs. K. Claiborn, E. Walker, B. Oleson, P. Arvan, J. Hansen, and members of the Soleimanpour laboratory for helpful advice. V.S. conceived, designed and performed experiments, interpreted results, drafted and reviewed the manuscript. G.L.P, V.P., B.T., L.C., M.A.G., J.Z., T.S., E.R., and B.C. designed and performed experiments and interpreted results. J.A.C., T.M-P., and L.S.S. designed studies, interpreted results and reviewed the manuscript. S.A.S. conceived and designed the studies, interpreted results, edited and reviewed the manuscript. All authors declare that they have no conflicts of interest.

## References

1. Donath MY, Dinarello CA, and Mandrup-Poulsen T. Targeting innate immune mediators in type 1 and type 2 diabetes. Nat Rev Immunol. 2019;19(12):734–46.

2. Imai Y, Dobrian AD, Morris MA, and Nadler JL. Islet inflammation: a unifying target for diabetes treatment? Trends Endocrinol Metab. 2013;24(7):351–60.

3. Padgett LE, Broniowska KA, Hansen PA, Corbett JA, and Tse HM. The role of reactive oxygen species and proinflammatory cytokines in type 1 diabetes pathogenesis. Ann N Y Acad Sci. 2013;1281:16–35.

4. Cerf ME. Beta cell dysfunction and insulin resistance. Front Endocrinol (Lausanne). 2013;4:37.

5. Cnop M, Welsh N, Jonas JC, Jorns A, Lenzen S, and Eizirik DL. Mechanisms of pancreatic beta-cell death in type 1 and type 2 diabetes: many differences, few similarities. Diabetes. 2005;54 Suppl 2:S97–107.

6. Hohmeier HE, Tran VV, Chen G, Gasa R, and Newgard CB. Inflammatory mechanisms in diabetes: lessons from the beta-cell. Int J Obes Relat Metab Disord. 2003;27 Suppl 3:S12–6.

7. Chen J, Stimpson SE, Fernandez-Bueno GA, and Mathews CE. Mitochondrial Reactive Oxygen Species and Type 1 Diabetes. Antioxid Redox Signal. 2018;29(14):1361–72.

8. Gustafsson AB, and Dorn GW, 2nd. Evolving and Expanding the Roles of Mitophagy as a Homeostatic and Pathogenic Process. Physiol Rev. 2019;99(1):853–92.

9. Kaufman BA, Li C, and Soleimanpour SA. Mitochondrial regulation of beta-cell function: maintaining the momentum for insulin release. Molecular aspects of medicine. 2015;42:91–104.

10. Sprenger HG, and Langer T. The Good and the Bad of Mitochondrial Breakups. Trends Cell Biol. 2019;29(11):888–900.

11. Pearson G, Chai B, Vozheiko T, Liu X, Kandarpa M, Piper RC, et al. Clec16a, Nrdp1, and USP8 Form a Ubiquitin-Dependent Tripartite Complex That Regulates beta-Cell Mitophagy. Diabetes. 2018;67(2):265–77.

12. Pearson G, and Soleimanpour SA. A ubiquitin-dependent mitophagy complex maintains mitochondrial function and insulin secretion in beta cells. Autophagy. 2018;14(7):1160–1.

13. Soleimanpour SA, Gupta A, Bakay M, Ferrari AM, Groff DN, Fadista J, et al. The diabetes susceptibility gene Clec16a regulates mitophagy. Cell. 2014;157(7):1577–90.

14. Dupuis J, Langenberg C, Prokopenko I, Saxena R, Soranzo N, Jackson AU, et al. New genetic loci implicated in fasting glucose homeostasis and their impact on type 2 diabetes risk. Nat Genet. 2010;42(2):105–16.

15. Chen L, Liu C, Gao J, Xie Z, Chan LWC, Keating DJ, et al. Inhibition of Miro1 disturbs mitophagy and pancreatic beta-cell function interfering insulin release via IRS-Akt-Foxo1 in diabetes. Oncotarget. 2017;8(53):90693–705.

16. Thorens B, Tarussio D, Maestro MA, Rovira M, Heikkila E, and Ferrer J. Ins1(Cre) knock-in mice for beta cell-specific gene recombination. Diabetologia. 2015;58(3):558–65.

17. Sun N, Yun J, Liu J, Malide D, Liu C, Rovira, II, et al. Measuring In Vivo Mitophagy. Mol Cell. 2015;60(4):685–96.

18. Laubach VE, Shesely EG, Smithies O, and Sherman PA. Mice lacking inducible nitric oxide synthase are not resistant to lipopolysaccharide-induced death. Proc Natl Acad Sci U S A. 1995;92(23):10688–92.

19. Gu G, Dubauskaite J, and Melton DA. Direct evidence for the pancreatic lineage: NGN3+ cells are islet progenitors and are distinct from duct progenitors. Development. 2002;129(10):2447–57.

20. Hodson DJ, Tarasov AI, Gimeno Brias S, Mitchell RK, Johnston NR, Haghollahi S, et al. Incretin-modulated beta cell energetics in intact islets of Langerhans. Mol Endocrinol. 2014;28(6):860–71.

21. Corsa CAS, Pearson GL, Renberg A, Askar MM, Vozheiko T, MacDougald OA, et al. The E3 ubiquitin ligase parkin is dispensable for metabolic homeostasis in murine pancreatic beta cells and adipocytes. J Biol Chem. 2019;294(18):7296–307.

22. Oleson BJ, Broniowska KA, Schreiber KH, Tarakanova VL, and Corbett JA. Nitric oxide induces ataxia telangiectasia mutated (ATM) protein-dependent gammaH2AX protein formation in pancreatic beta cells. J Biol Chem. 2014;289(16):11454–64.

23. Hansen JB, Tonnesen MF, Madsen AN, Hagedorn PH, Friberg J, Grunnet LG, et al. Divalent metal transporter 1 regulates iron-mediated ROS and pancreatic beta cell fate in response to cytokines. Cell Metab. 2012;16(4):449–61.

24. Sun N, Malide D, Liu J, Rovira, II, Combs CA, and Finkel T. A fluorescence-based imaging method to measure in vitro and in vivo mitophagy using mt-Keima. Nat Protoc. 2017;12(8):1576–87.

25. Nezich CL, Wang C, Fogel AI, and Youle RJ. MiT/TFE transcription factors are activated during mitophagy downstream of Parkin and Atg5. J Cell Biol. 2015;210(3):435–50.

26. Rojansky R, Cha MY, and Chan DC. Elimination of paternal mitochondria in mouse embryos occurs through autophagic degradation dependent on PARKIN and MUL1. Elife. 2016;5.

27. Merrins MJ, Poudel C, McKenna JP, Ha J, Sherman A, Bertram R, et al. Phase Analysis of Metabolic Oscillations and Membrane Potential in Pancreatic Islet beta-Cells. Biophys J. 2016;110(3):691–9.

28. Berg J, Hung YP, and Yellen G. A genetically encoded fluorescent reporter of ATP:ADP ratio. Nat Methods. 2009;6(2):161–6.

29. Bříza T, Rimpelová S, Králová J, Záruba K, Kejík Z, Ruml T, et al. Pentamethinium fluorescent probes: The impact of molecular structure on photophysical properties and subcellular localization. Dyes Pigm. 2014;107:51–9.

30. Jayaraman S. A novel method for the detection of viable human pancreatic beta cells by flow cytometry using fluorophores that selectively detect labile zinc, mitochondrial membrane potential and protein thiols. Cytometry A. 2008;73(7):615–25.

31. Chaudhry A, Shi R, and Luciani DS. A pipeline for multidimensional confocal analysis of mitochondrial morphology, function and dynamics in pancreatic beta-cells. Am J Physiol Endocrinol Metab. 2019.

32. Vowinckel J, Hartl J, Butler R, and Ralser M. MitoLoc: A method for the simultaneous quantification of mitochondrial network morphology and membrane potential in single cells. Mitochondrion. 2015;24:77–86.

33. Jin SM, and Youle RJ. PINK1- and Parkin-mediated mitophagy at a glance. J Cell Sci. 2012;125(Pt 4):795–9.

34. Scarim AL, Heitmeier MR, and Corbett JA. Irreversible inhibition of metabolic function and islet destruction after a 36-hour exposure to interleukin-1beta. Endocrinology. 1997;138(12):5301–7.

35. Corbett JA, Wang JL, Sweetland MA, Lancaster JR, Jr., and McDaniel ML. Interleukin 1 beta induces the formation of nitric oxide by beta-cells purified from rodent islets of Langerhans. Evidence for the beta-cell as a source and site of action of nitric oxide. J Clin Invest. 1992;90(6):2384–91.

36. Welsh N, Eizirik DL, Bendtzen K, and Sandler S. Interleukin-1 beta-induced nitric oxide production in isolated rat pancreatic islets requires gene transcription and may lead to inhibition of the Krebs cycle enzyme aconitase. Endocrinology. 1991;129(6):3167–73.

37. McArdle F, Pattwell DM, Vasilaki A, McArdle A, and Jackson MJ. Intracellular generation of reactive oxygen species by contracting skeletal muscle cells. Free Radic Biol Med. 2005;39(5):651–7.

38. Arimura K, Egashira K, Nakamura R, Ide T, Tsutsui H, Shimokawa H, et al. Increased inactivation of nitric oxide is involved in coronary endothelial dysfunction in heart failure. Am J Physiol Heart Circ Physiol. 2001;280(1):H68–75.

39. Guzik TJ, West NE, Pillai R, Taggart DP, and Channon KM. Nitric oxide modulates superoxide release and peroxynitrite formation in human blood vessels. Hypertension. 2002;39(6):1088–94.

40. Marasco MR, Conteh AM, Reissaus CA, Cupit JEt, Appleman EM, Mirmira RG, et al. Interleukin-6 Reduces beta-Cell Oxidative Stress by Linking Autophagy With the Antioxidant Response. Diabetes. 2018;67(8):1576–88.

41. Raza H, Prabu SK, John A, and Avadhani NG. Impaired mitochondrial respiratory functions and oxidative stress in streptozotocin-induced diabetic rats. Int J Mol Sci. 2011;12(5):3133–47.

42. Rossini AA, Like AA, Chick WL, Appel MC, and Cahill GF, Jr. Studies of streptozotocin-induced insulitis and diabetes. Proc Natl Acad Sci U S A. 1977;74(6):2485–9.

43. Twig G, Elorza A, Molina AJ, Mohamed H, Wikstrom JD, Walzer G, et al. Fission and selective fusion govern mitochondrial segregation and elimination by autophagy. EMBO J. 2008;27(2):433–46.

44. Basiorka AA, McGraw KL, De Ceuninck L, Griner LN, Zhang L, Clark JA, et al. Lenalidomide Stabilizes the Erythropoietin Receptor by Inhibiting the E3 Ubiquitin Ligase RNF41. Cancer Res. 2016;76(12):3531–40.

45. Gingerich MA, Sidarala V, and Soleimanpour SA. Clarifying the function of genes at the chromosome 16p13 locus in type 1 diabetes: CLEC16A and DEXI. Genes Immun. 2020;21(2):79–82.

46. Hakonarson H, Grant SF, Bradfield JP, Marchand L, Kim CE, Glessner JT, et al. A genome-wide association study identifies KIAA0350 as a type 1 diabetes gene. Nature. 2007;448(7153):591–4.

47. Nieves-Bonilla JM, Kiaf B, Schuster C, and Kissler S. The type 1 diabetes candidate gene Dexi does not affect disease risk in the nonobese diabetic mouse model. Genes Immun. 2019.

48. Dooley J, Tian L, Schonefeldt S, Delghingaro-Augusto V, Garcia-Perez JE, Pasciuto E, et al. Genetic predisposition for beta cell fragility underlies type 1 and type 2 diabetes. Nat Genet. 2016;48(5):519–27.

49. Peters L, Posgai A, and Brusko TM. Islet-immune interactions in type 1 diabetes: the nexus of beta cell destruction. Clin Exp Immunol. 2019;198(3):326–40.

50. Li J, Jorgensen SF, Maggadottir SM, Bakay M, Warnatz K, Glessner J, et al. Association of CLEC16A with human common variable immunodeficiency disorder and role in murine B cells. Nat Commun. 2015;6:6804.

51. Pandey R, Bakay M, Hain HS, Strenkowski B, Elsaqa BZB, Roizen JD, et al. CLEC16A regulates splenocyte and NK cell function in part through MEK signaling. PLoS One. 2018;13(9):e0203952.

52. Pandey R, Bakay M, Hain HS, Strenkowski B, Yermakova A, Kushner JA, et al. The Autoimmune Disorder Susceptibility Gene CLEC16A Restrains NK Cell Function in YTS NK Cell Line and Clec16a Knockout Mice. Front Immunol. 2019;10:68.

53. Redmann V, Lamb CA, Hwang S, Orchard RC, Kim S, Razi M, et al. Clec16a is Critical for Autolysosome Function and Purkinje Cell Survival. Sci Rep. 2016;6:23326.

54. Schuster C, Gerold KD, Schober K, Probst L, Boerner K, Kim MJ, et al. The Autoimmunity-Associated Gene CLEC16A Modulates Thymic Epithelial Cell Autophagy and Alters T Cell Selection. Immunity. 2015;42(5):942–52.

55. van Luijn MM, Kreft KL, Jongsma ML, Mes SW, Wierenga-Wolf AF, van Meurs M, et al. Multiple sclerosis-associated CLEC16A controls HLA class II expression via late endosome biogenesis. Brain. 2015;138(Pt 6):1531–47.

56. Chen J, Chernatynskaya AV, Li JW, Kimbrell MR, Cassidy RJ, Perry DJ, et al. T cells display mitochondria hyperpolarization in human type 1 diabetes. Sci Rep. 2017;7(1):10835.

57. Zhong Z, Umemura A, Sanchez-Lopez E, Liang S, Shalapour S, Wong J, et al. NF-kappaB Restricts Inflammasome Activation via Elimination of Damaged Mitochondria. Cell. 2016;164(5):896–910.

58. Zhou R, Yazdi AS, Menu P, and Tschopp J. A role for mitochondria in NLRP3 inflammasome activation. Nature. 2011;469(7329):221–5.

59. Glick D, Barth S, and Macleod KF. Autophagy: cellular and molecular mechanisms. J Pathol. 2010;221(1):3–12.

60. Jung HS, Chung KW, Won Kim J, Kim J, Komatsu M, Tanaka K, et al. Loss of autophagy diminishes pancreatic beta cell mass and function with resultant hyperglycemia. Cell Metab. 2008;8(4):318–24.

61. Marasco MR, and Linnemann AK. beta-Cell Autophagy in Diabetes Pathogenesis. Endocrinology. 2018;159(5):2127–41.

62. Nakatogawa H, Suzuki K, Kamada Y, and Ohsumi Y. Dynamics and diversity in autophagy mechanisms: lessons from yeast. Nat Rev Mol Cell Biol. 2009;10(7):458–67.

63. Petcherski A, Trudeau KM, Wolf DM, Segawa M, Lee J, Taddeo EP, et al. Elamipretide Promotes Mitophagosome Formation and Prevents Its Reduction Induced by Nutrient Excess in INS1 beta-cells. J Mol Biol. 2018;430(24):4823–33.

64. Cerqueira FM, Kozer N, Petcherski A, Baranovski BM, Wolf D, Assali EA, et al. MitoTimer-based high-content screen identifies two chemically-related benzothiophene derivatives that enhance basal mitophagy. Biochem J. 2020;477(2):461–75.

65. Ryu D, Mouchiroud L, Andreux PA, Katsyuba E, Moullan N, Nicolet-Dit-Felix AA, et al. Urolithin A induces mitophagy and prolongs lifespan in C. elegans and increases muscle function in rodents. Nat Med. 2016;22(8):879–88.

66. Allen GF, Toth R, James J, and Ganley IG. Loss of iron triggers PINK1/Parkin-independent mitophagy. EMBO Rep. 2013;14(12):1127–35.

67. Andreux PA, Blanco-Bose W, Ryu D, Burdet F, Ibberson M, Aebischer P, et al. The mitophagy activator urolithin A is safe and induces a molecular signature of improved mitochondrial and cellular health in humans. Nature Metabolism. 2019;1(6):595–603.

