## Supplementary Figures and Legends for "Mitophagy protects beta cells from inflammatory damage in diabetes"

**Supplementary Figure Legends**

**Figure S1. Evaluation of mitophagy and bulk macroautophagy following mitochondrial damage.**

**(A)** Mfn1 and Mfn2 expression by WB in Min6 β-cells treated with 5μM FCCP for 6hrs or 1μM valinomycin for indicated time course. **(B)** LC3 expression by WB in primary mouse islets treated with cytokines for 24hrs. n=3/group for all studies.

**Figure S2. Mitochondrial fragmentation in mitophagy-deficient human β-cells following pro-inflammatory cytokines.**

Human β-cell mitochondrial network analysis of confocal immunofluorescence Z-stack images stained for SDHA from studies depicted in Figure 5F by MitoAnalyzer. n=3/group (75-110 β-cells from each donor per condition were quantified). * $p<0.05$ vs. Veh + PBS. $ $p<0.005$ vs. Veh + Cyt.

**Figure S3. Mitophagy deficiency does not induce differences in inflammatory mediators following cytokine exposure.**

**(A)** Clec16a mRNA expression (normalized to Hprt) by qRT-PCR from RNA isolated from shNT or shClec16a-expressing Min6 β-cells treated in the presence/absence of cytokines for 24hrs. * $p<0.05$ vs. shNT + PBS. $ $p<0.05$ vs. shNT + Cyt. **(B)** Nitrite release (normalized to total protein content) in cell culture supernatants of NT or Clec16a-specific shRNA expressing Min6 β-cells treated with/without cytokines for 24hrs. * $p<0.05$ vs. shNT + PBS; $ $p<0.05$ vs. shNT + PBS. **(C)** Total cellular reactive oxygen species (normalized to total protein levels) in NT or Clec16a-specific shRNA expressing Min6 β-cells incubated with/without 500μM tiron for 48hrs and treated with/without cytokines for the final 24hrs. * $p<0.05$ vs. shNT + PBS; $ $p<0.05$ vs. shNT + Cyt. **(D)** qRT-PCR of selected NFκB transcriptional targets (normalized to Hprt) from RNA isolated from shNT or shClec16a-expressing Min6 β-cells in the presence/absence of cytokines for 24hrs. *p<0.05 vs. shNT + PBS. **(E)** Cellular labile iron pools (LIP; measured in arbitrary units) in shNT or shClec16a-expressing Min6 β-cells treated in the presence/absence of cytokines for 24hrs. *p< 0.05, **p< 0.01. **(F)** NOS2 mRNA expression (normalized to Cyclophilin A; CYPA) by qRT-PCR from RNA isolated from human islets treated with/without 10 μM lenalidomide for 48hrs and then treated with/without cytokines for the final 24hrs. * p<0.05 vs. Veh + PBS. **(G)** qRT-PCR of other selected NFκB transcriptional targets (normalized to CYPA) from RNA isolated from human islets treated with/without 10 μM lenalidomide for 48hrs and then exposed to cytokines or vehicle for the final 24hrs. * p<0.05 vs. Veh + PBS. n=3-10/group for all studies.

**Figure S4. Clec16a overexpression protects against cytokine toxicity.**

**(A)** Immunofluorescence image of human islets transduced with empty vector control (Ad.EV) or Clec16a-overexpressing (Ad.Clec16a) adenoviral particles stained for insulin (red), GFP (green), and DAPI (blue). β-cells were identified by insulin immunostaining, and virally transduced cells were identified by GFP immunostaining. **(B)** Flag-Clec16a expression by WB performed in Min6 β-cells, 72hrs after overnight transduction with Ad.GFP or Ad.Clec16a adenoviral particles. Cyclophilin B serves as a loading control. (**C)** Immunofluorescence image of human islets, transduced with empty vector control (Ad.EV) or Clec16a-overexpressing (Ad.Clec16a) adenoviral particles, and exposed to cytokines or vehicle for 24hrs stained for TUNEL (red), GFP (green), and DAPI (blue). Transduced cells were identified by GFP immunostaining. **(D)** Cleaved caspase 3 and Flag expression by WB in empty vector (EV) or Clec16a-Flag expressing Min6 β-cells treated with cytokines for 6hrs. **(E)** Cleaved caspase 3 densitometry (normalized to Cyclophilin B) from studies depicted in Figure S4D. n=3/group. *p<0.05 vs. EV + PBS; $ p<0.05 vs. EV + Cyt.

A)

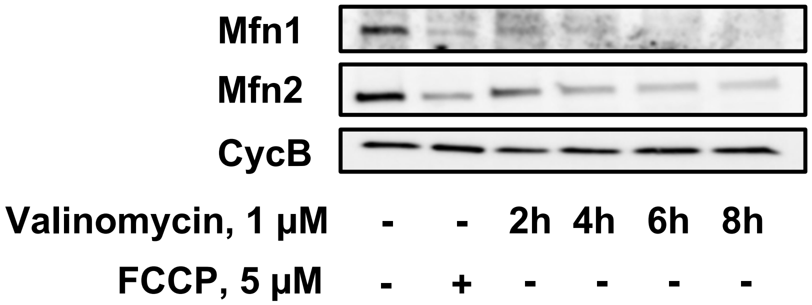
Mfn1
Mfn2
CycB
Valinomycin, 1 µM - - 2h 4h 6h 8h
FCCP, 5 µM - + - - - -

B)

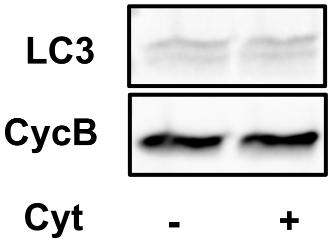
LC3
CycB
Cyt - +

### Figure S2

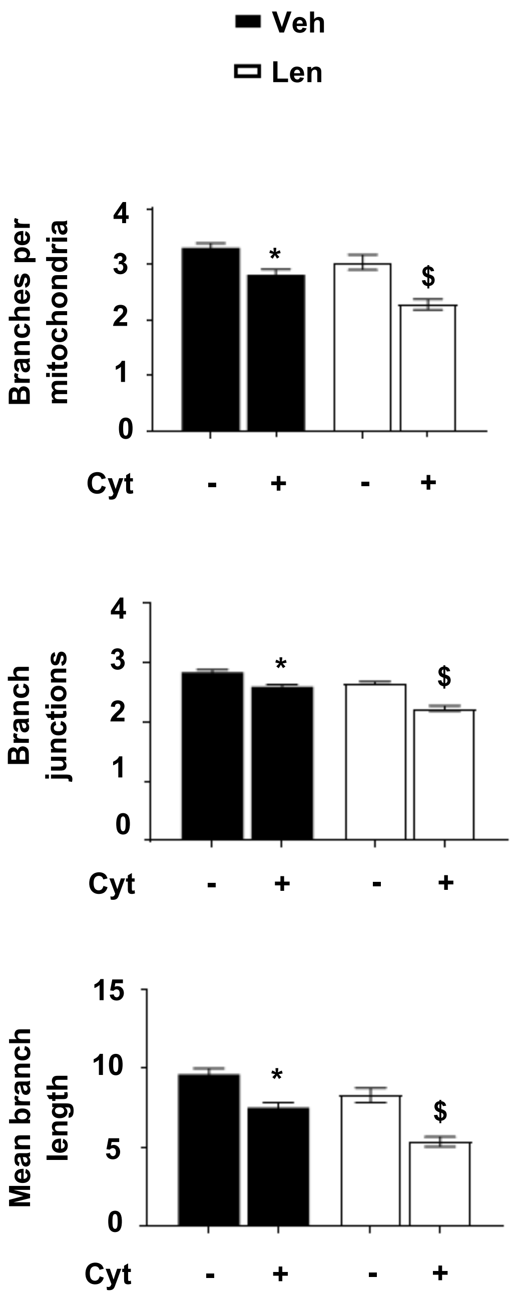

Veh
Len
Branches per mitochondria
4
3
2
1
0
*
$
Cyt
-
+
-
+
Branch junctions
4
3
2
1
0
*
$
Cyt
-
+
-
+
Mean branch length
15
10
5
0
*
$
Cyt
-
+
-
+

Figure S3

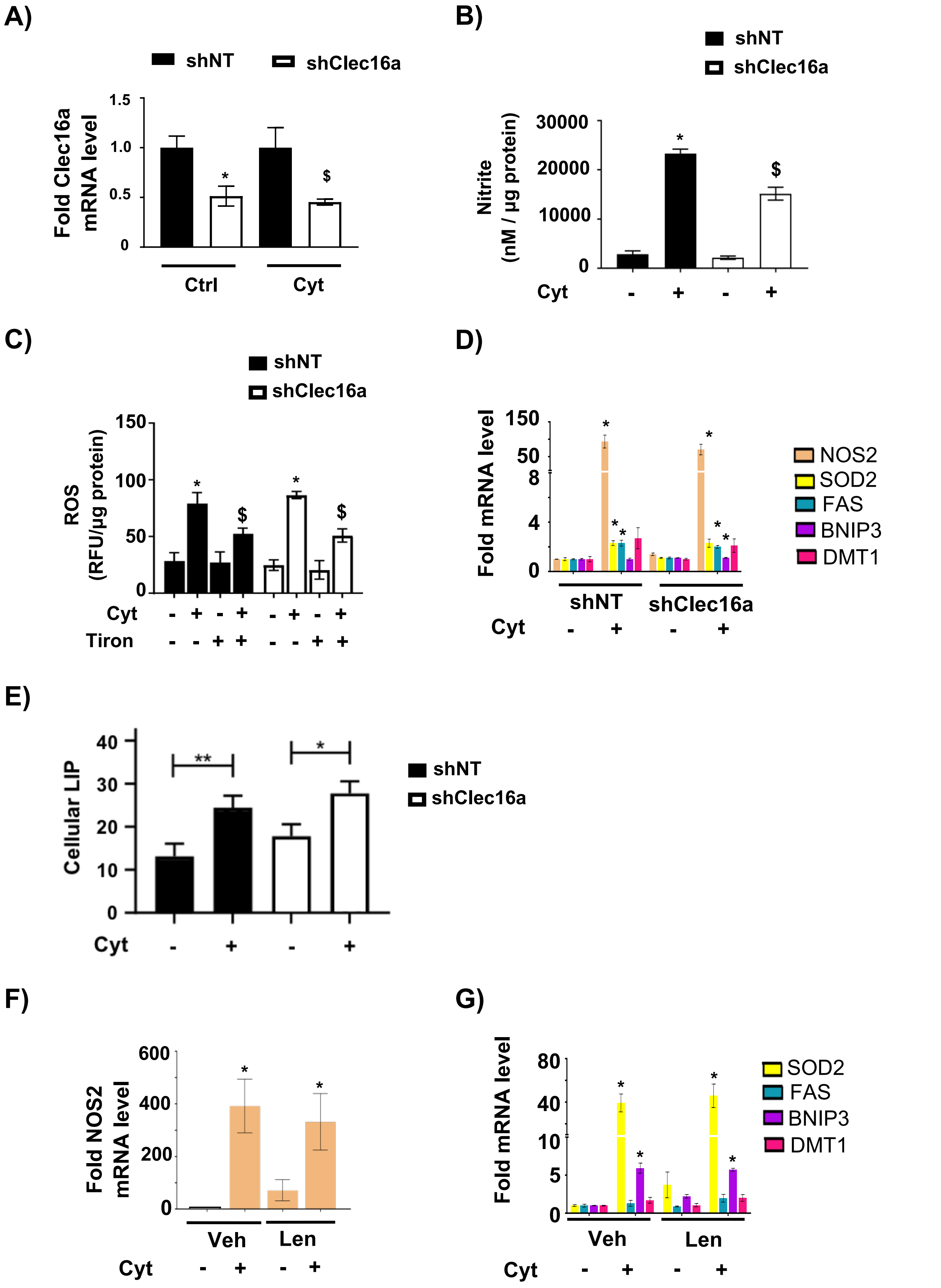
A)
shNT
shClec16a
Fold Clec16a mRNA level
1.5
1.0
0.5
0
*
$
Ctrl
Cyt
B)
shNT
shClec16a
Nitrite (nM / µg protein)
30000
20000
10000
0
*
$
Cyt - + - +
C)
shNT
shClec16a
ROS (RFU/µg protein)
150
100
50
0
*
$
*
$
Cyt - + - + - + - +
Tiron - - + + - - + +
D)
Fold mRNA level
150
50
8
4
0
*
*
*
*
*
*
NOS2
SOD2
FAS
BNIP3
DMT1
shNT
shClec16a
Cyt - + - +
E)
Cellular LIP
40
30
20
10
0
**
*
shNT
shClec16a
Cyt - + - +
F)
Fold NOS2 mRNA level
600
400
200
0
*
*
Veh
Len
Cyt - + - +
G)
Fold mRNA level
80
40
10
5
0
*
*
*
*
SOD2
FAS
BNIP3
DMT1
Veh
Len
Cyt - + - +

### Figure S4

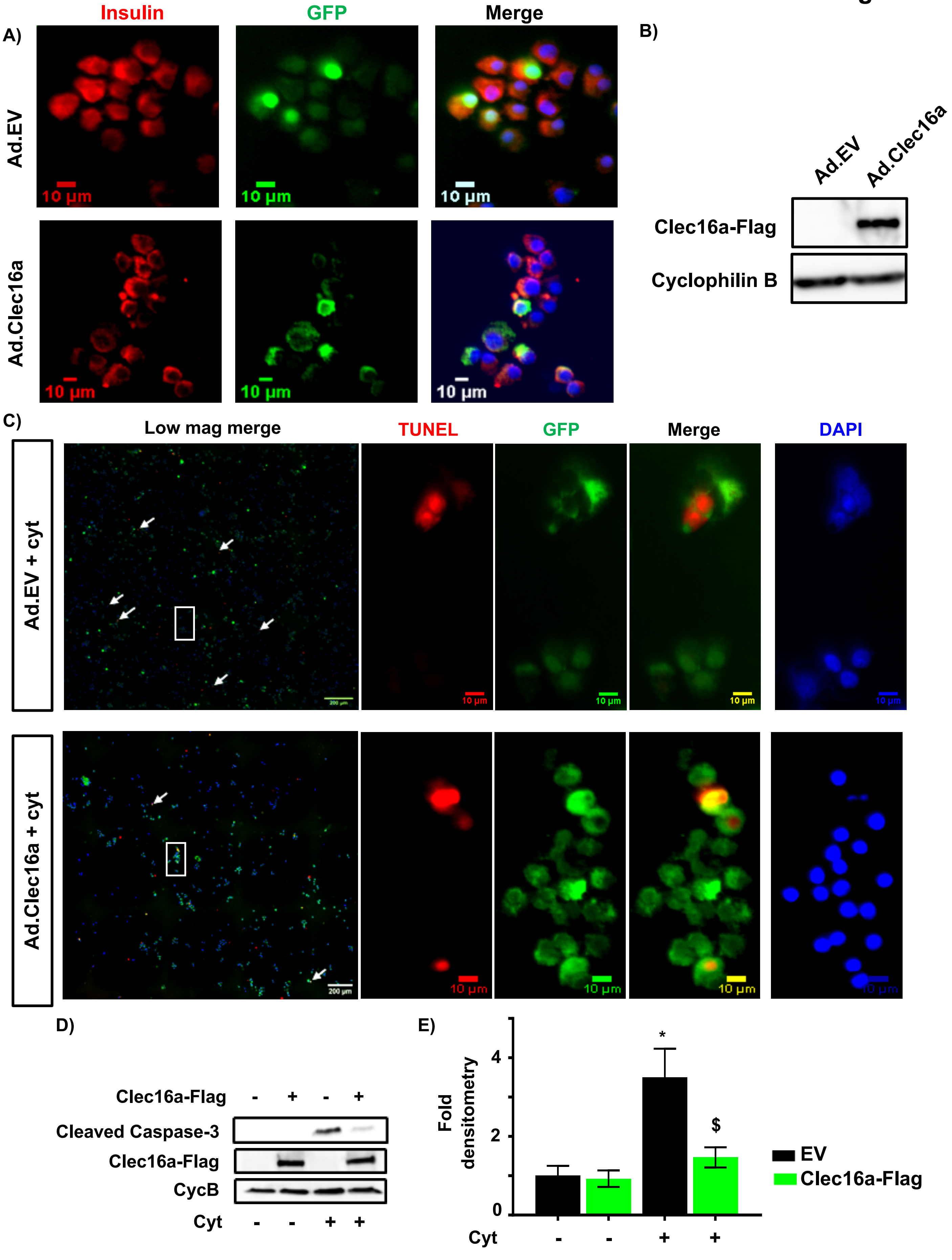
A)
Insulin
GFP
Merge
Ad.EV
Ad.Clec16a
10 µm
B)
Ad.EV
Ad.Clec16a
Clec16a-Flag
Cyclophilin B
C)
Low mag merge
TUNEL
GFP
Merge
DAPI
Ad.EV + cyt
Ad.Clec16a + cyt
10 µm
200 µm
D)
Clec16a-Flag - + - +
Cleaved Caspase-3
Clec16a-Flag
CycB
Cyt - - + +
E)
Fold densitometry
0
2
4
*
$
EV
Clec16a-Flag
Cyt - - + +
