## Supplementary Table 2 for "Mitophagy protects beta cells from inflammatory damage in diabetes"

**Supplementary Table 2. Commercially available antibodies used within this study.**

| Antibody | Company | Catalog # |
|---|---|---|
| β-Actin | ThermoFisher | MA5-15739 (BA3R) |
| Cleaved Caspase 3 | Cell Signaling | 9964 (5A1E) |
| Cyclophilin B | ThermoFisher | PA1-027A |
| Flag | Sigma | F1804 (clone M2) |
| GFP | Abcam | ab6673 |
| Glucagon | Santa Cruz | sc-13091 |
| Insulin | Dako | A0564 |
| Lamp1 | Developmental Studies Hybridoma Bank | 1D4B |
| LC3 | Sigma | L8918 |
| Mfn1 | Abcam | ab126575 (11E91H12) |
| Mfn2 | Abcam | ab56889 (6A8) |
| NOS2 | Cayman Chemical | 160862 |
| Parkin | EMD Millipore | 05-882 (PRK8) |
| Pdx1 | Abcam | ab47383 |
| SDHA | Abcam | ab14715 (2E3GC12FB2AE2) |
