## Supplementary Table 3 for "Mitophagy protects beta cells from inflammatory damage in diabetes"

**Supplementary Table 3. qRT-PCR primer sequences.**

| Gene | Target species | Fwd | Rev |
|---|---|---|---|
| Clec16a | Mouse | 5'-TGCAGCTGCTACAGACCTTGA-3' | 5'-ATACGCCATGATCTCCTCGTC-3' |
| Nos2 | Mouse | 5'-ACTGGGACAGCACAGAATGTTCC-3' | 5'-CCAAATGTGCTTGTCACCACCAG-3' |
| Sod2 | Mouse | 5'-TACAACTCAGGTCGCTCTTCAGC-3' | 5'-AGCCTCCAGCAACTCTCCTTT-3' |
| Fas | Mouse | 5'-TTAAAGCTGAGGAGGCGGGTT-3' | 5'-CTCAGCCTAGTTTTCAGGTTGGC-3' |
| Bnip3 | Mouse | 5'-GCTTGGGGATCTACATTGGAAGG-3' | 5'-GTGCAAACACCCAAGGACCAT-3' |
| Dmt1 | Mouse | 5'-GCATTGGGTCTGTCTTTCCTG-3' | 5'-TGGACACCACTGAGTCAGCAT-3' |
| Hprt | Mouse | 5'-TGCTCGAGATGTCATGAAGGA-3' | 5'-CCAGCAGGTCAGCAAAGAACT-3' |
| NOS2 | Human | 5'-TGCCCTGGCAATGGAGAGAAA-3' | 5'-GCCAAACACAGCGTACCTGAA-3' |
| SOD2 | Human | 5'-TAGCTCTTCAGCCTGCACTGA-3' | 5'-AGCAACTCCCCTTTGGGTTCT-3' |
| FAS | Human | 5'-TGTCCTCCAGGTGAAAGGAAAGC-3' | 5'-TGTACTCCTTCCCTTCTTGGCAG-3' |
| BNIP3 | Human | 5'-CTCTGCTGCTCTCTCATTTGCTG-3' | 5'-AAAGGTGCTGGTGGAGGTTGT-3' |
| DMT1 | Human | 5'-ACCAACGAGCAGGTGGTTGAA-3' | 5'-AGTGCAGCAGGCCCAAAGTAA-3' |
| CYPA | Human | 5'-GCGTCTCCTTTGAGCTGTTTGCA-3' | 5'-CCACCCTGACACATAAACCCTGGAA-3' |
